# GLDADec: marker-gene guided LDA modelling for bulk gene expression deconvolution

**DOI:** 10.1101/2024.01.08.574749

**Authors:** Iori Azuma, Tadahaya Mizuno, Hiroyuki Kusuhara

**Author notes:** Corresponding author: Tadahaya Mizuno; Graduate School of Pharmaceutical Sciences, the University of Tokyo, Bunkyo-ku, Tokyo, 113-0033, Japan. These authors contributed equally (Iori Azuma, Tadahaya Mizuno).

## Abstract

Inferring cell type proportions from bulk transcriptome data is crucial in immunology and oncology. Here, we introduce GLDADec (Guided LDA Deconvolution), a bulk deconvolution method that guides topics using cell type-specific marker gene names to estimate topic distributions for each sample. Through benchmarking using blood-derived datasets, we demonstrate its high estimation performance and robustness. Moreover, we apply GLDADec to heterogeneous tissue bulk data and perform comprehensive cell type analysis in a data-driven manner. We show that GLDADec outperforms existing methods in estimation performance and evaluate its biological interpretability by examining enrichment of biological processes for topics. Finally, we apply GLDADec to TCGA tumor samples, enabling subtype stratification and survival analysis based on estimated cell type proportions, thus proving its practical utility in clinical settings. This approach, utilizing marker gene names as partial prior information, can be applied to various scenarios for bulk data deconvolution. GLDADec is available as an open-source Python package at https://github.com/mizuno-group/GLDADec.

## Introduction

Quantification of the proportion of cell types in a tissue sample and understanding the contribution of individual cell types to the physiological state, such as immune responses associated with perturbation or evaluation of cancer tumor samples with cell proliferation, is of utmost importance [1,2]. Flow cytometry, a typical experimental approach for quantifying compositional proportions of cell types, is widely used in molecular biology and immunology. However, its application is limited to fresh organs, human tissue specimens are challenging to analyze, and there is limited knowledge on the aggregation of flow cytometry data, making it difficult to utilize legacy data.

The advancement of high-throughput sequencing technology has led to an abundance of stored transcriptome data [3,4]. The bulk transcriptome measures the accumulation of gene expression levels derived from various cell types and can be applied for extensive analyses using several well-established databases [3–5]. Although databases for single-cell sequencing technology, a recent innovation, have also been developed, its high cost and sparce nature make it challenging to conduct large-scale data analyses [6]. Hence, it is useful to establish a method to estimate the proportion of constitutive cells from the bulk transcriptome.

Deconvolution is a computational method that can be used to estimate the proportion of immune cells in a sample using transcriptome data. In recent years, several deconvolution methods have been proposed to infer cell type proportions from bulk expression data [7–14]. These methods can be categorized into two main groups: reference-free methods and reference-based methods [15,16] Reference-free methods estimate cell type proportions solely based on the samples to be analyzed, making them less sensitive to external information that may cause confounding factors. This approach is reasonable for tissue data where the constituent cell types are not clearly defined, as the number of components can be estimated using likelihood and other factors [11,12]. However, discerning the inferred components and their associated cells poses a challenge, particularly in more detailed cell types, thereby rendering the downstream task less interpretable. In contrast, reference-based methods utilize cell type-specific gene expression profiles, called references, as prior information. Although there have been some notable successes [8,13,14], the performance of these methods depends on the quality of the reference data and the batch-to-batch differences between the data to be analyzed. As a result, reference-based methods are only applicable in specific scenarios where the major cellular components are well-defined and appropriate reference data are available. While they are effective for simulated datasets or samples with well-defined constituent cell types, such as blood, they may underestimate the impact of gene expression profiles derived from cells that are not assumed as the reference [17].

Latent Dirichlet Allocation (LDA) has been developed in the context of natural language documents [18,19]. The LDA model is designed to identify the topics that make up the content of documents, which is analogous to deconvolution, a method that extracts cell type-specific information from bulk transcriptome data. However, since LDA is an unsupervised learning method, it is classified as reference-free when simply applied to deconvolution, which presents a challenge in terms of cell identifiability. To address this issue, several approaches have been developed by incorporating prior information into the LDA algorithm and extending it to semi-supervised learning [12,20]. While these concepts are logical, they hinge on expression levels acquired from pure cell lines or single-cell RNA-Seq as prior knowledge, which is vulnerable to technical biases and imposes restrictive assumptions regarding distribution disparities from target bulk samples.

Here, we propose a novel Guided LDA Deconvolution (GLDADec) method, which utilizes marker gene names as partial prior information to estimate cell type proportions, thereby overcoming the challenges of conventional reference-based and reference-free methods simultaneously. This method employs a semi-supervised learning algorithm that combines cell-type marker genes with additional factors that may influence gene expression profiles to achieve a robust estimation of cell type proportions. Moreover, a median selection strategy is used to aggregate the outputs and achieve a more accurate estimation. We benchmarked GLDADec against existing methods using blood-derived samples with well-defined constituent cells, and it consistently outperformed the existing methods for multiple datasets. We also applied GLDADec to liver bulk RNA-Seq data from drug-induced liver injury models of mice and rats, demonstrating its usefulness for tissue data analysis. Our model, which considers additional topics, reflects the biological processes underlying the tissue and provides a robust estimation of the guided target cell types. Additionally, by collecting marker gene names in a data-driven manner and using them as prior information on comprehensive cell types, it is possible to estimate a wider range of cell types that are inaccessible by conventional methods. As a further demonstration, we applied GLDADec to human tumor samples, revealing new insights into cancer subtyping and clinical prognostic stratification. GLDADec is available as an open-source package at https://github.com/mizuno-group/GLDADec.

## Materials and Methods

### Overview of GLDADec

#### Latent Dirichlet Allocation (LDA) algorithm for deconvolution

The graphical model of the generation process is depicted in Figure 1. Each bulk gene expression sample indexed by m is the accumulation of a mixture of K cell types. It is assumed that the sample-topic distribution *θ*_m_ and topic-gene distribution φ_k_ are generated from the Dirichlet distribution as follows:

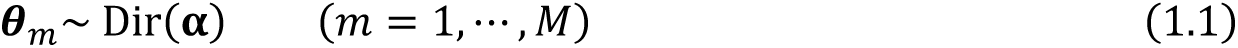

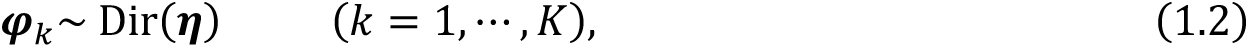

where α and η are hyperparameters. The latent topic *Z_m,n_* and gene *G_m,n_* are derived from multinomial distributions *Z_m,n_*∼Multi(*θ*_m_) and *G_m,n_*∼Multi(*φ*_m,n_), respectively. The joint probability is given by:

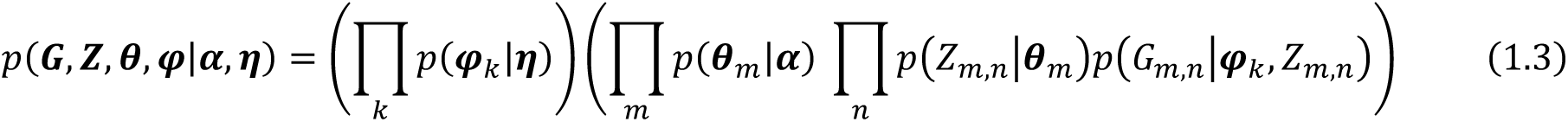

With respect to the observed gene expression levels, the probability distribution that generated them is approximated through collapsed Gibbs sampling [21]. Specifically, the posterior distribution is expressed as follows:

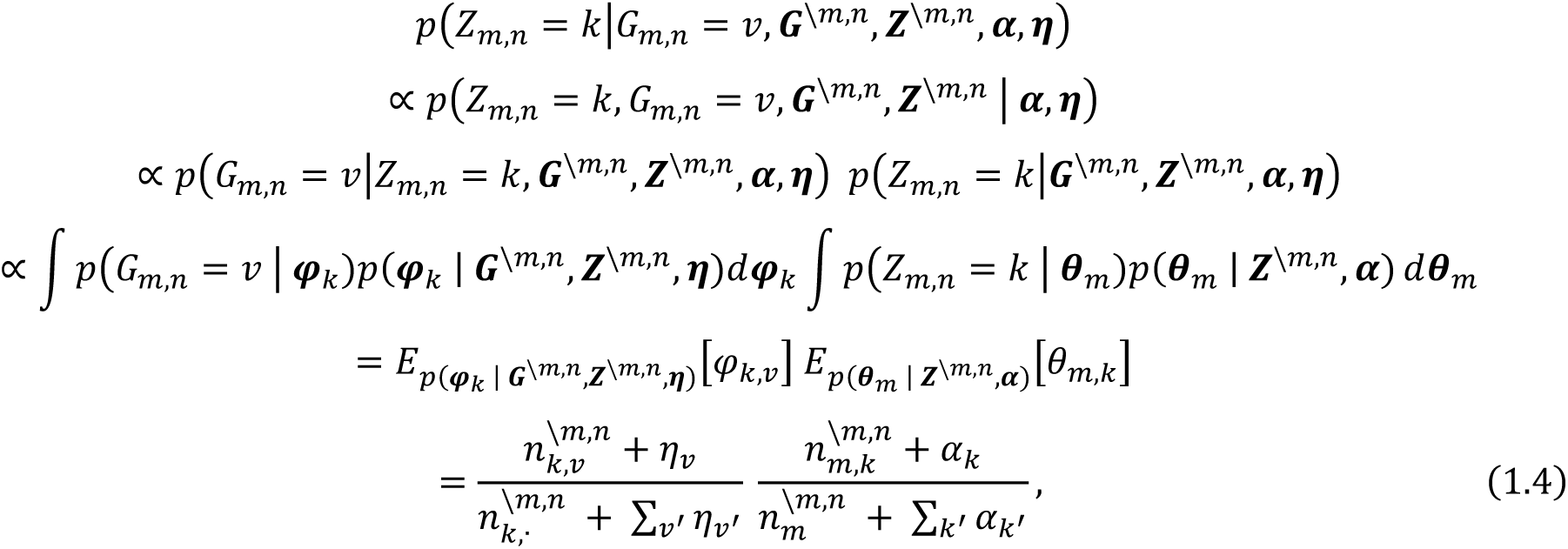

where *G^\m,n^* is the distribution without counting *G_m,n_* = v and *Z^\m,n^* is without counting *Z_m,n_* = *k*. Note that *n_k,v_^\m,n^* is the total number of counts allocated to topic *k* excluding the n^th^ gene *v* in sample *m*, and *n_m,k_^\m,n^* is the number of counts in sample *m* excluding the genes allocated to topic *k*, respectively. By utilizing these probability distributions, latent topics are sampled, and genes are assigned. The pseudo code for collapsed Gibbs sampling is described in Algorithm1.

The joint distribution of gene set *G* and topic set *Z* is given by integrating out *θ* and *φ* as follows:

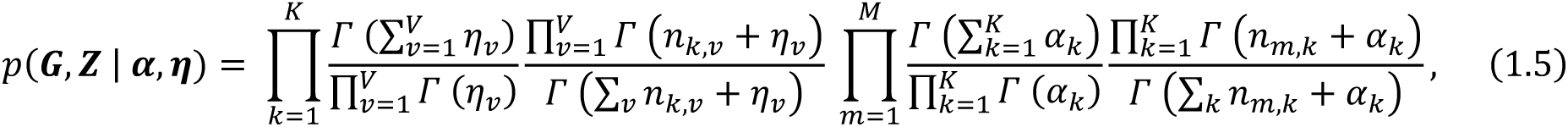

where 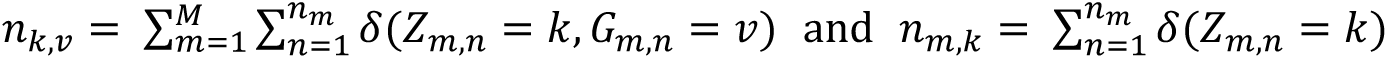.The proof of Equation 1.5 is provided in Supplementary Note S3. Then we have the log-likelihood at the sth step of the sampling process:

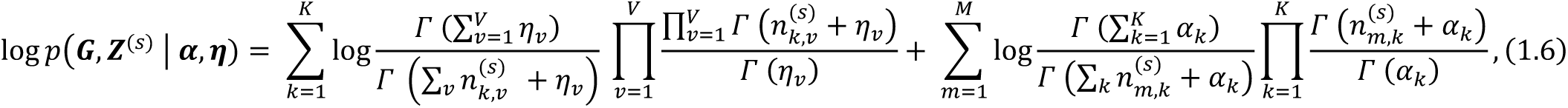

and the convergence during iteration is monitored with this.

The LDA algorithm provides the topic distribution for each sample **θ**_m_ and the gene distribution for each topic **φ**_k_ as output. In the context of cell-type deconvolution, the topics in the obtained **θ**_m_ correspond to the cell types that make up the sample.

**Fig. 1.**
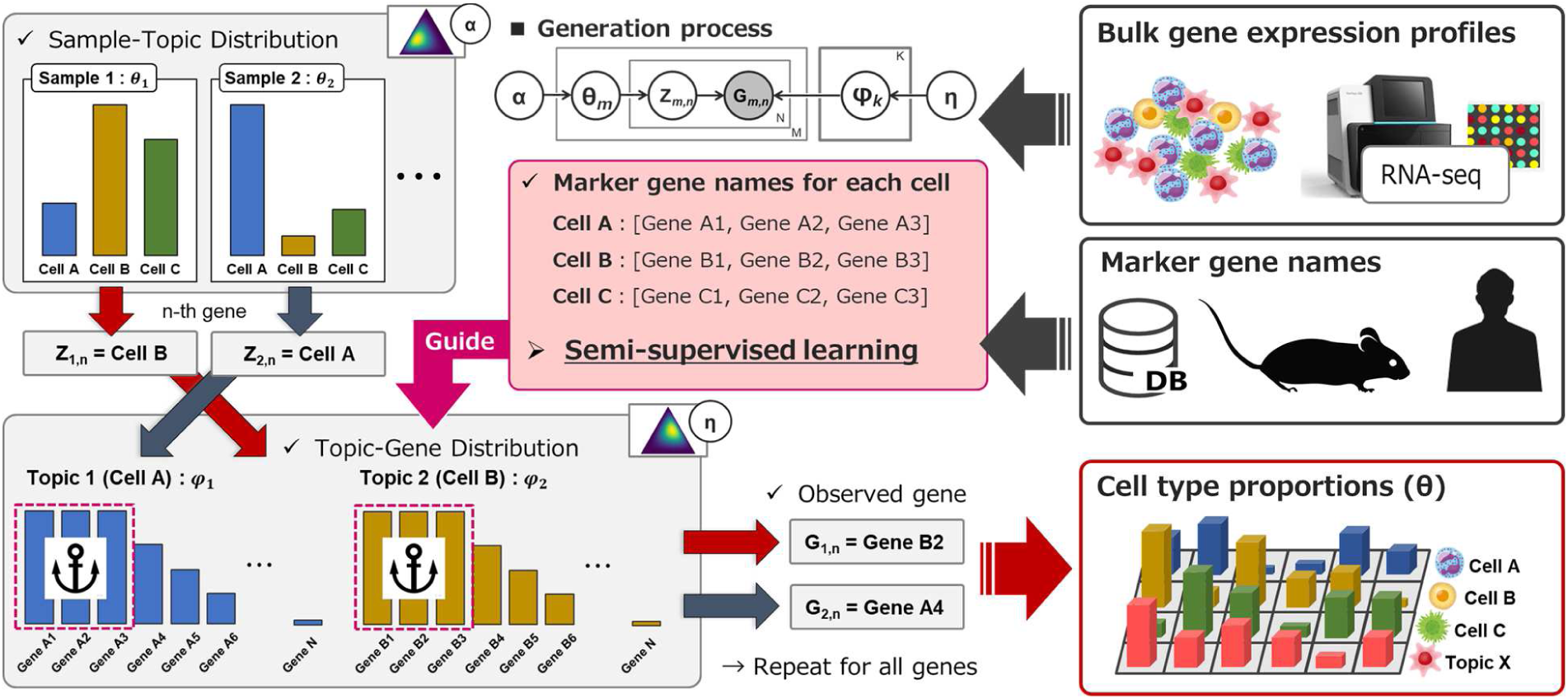
Overview of GLDADec. The observed gene expression profiles are considered as bag-of-words. We extend the standard LDA generation process to incorporate semi-supervised learning, where the gene names specific to each cell (topic) serve as partial prior information to guide the process. By running GLDADec, we can obtain θ, which reflects the cell type proportions in each sample.

#### Marker gene name based guided LDA algorithm

The LDA algorithm is an unsupervised learning algorithm that poses a challenge in identifying the cell type corresponding to a particular topic. To address this issue, guided LDA models have been proposed, which incorporate prior information to guide the topics and specify a direction towards the generation process of the LDA model [22,23]. In the context of deconvolution, the marker gene set of the cell assumed as topic *k*, ***S****_k_*, is used to guide the LDA algorithm as follows:

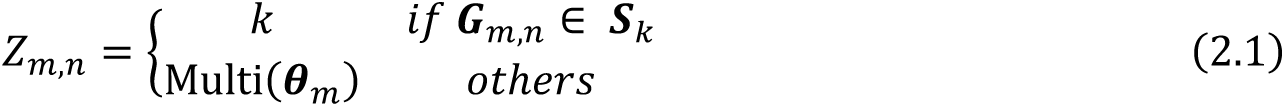

where the general concept of this process is to initialize the topic-gene distribution and to link topics to cells and allow identification of specific cell proportions. During the learning process, the contribution of marker genes to the corresponding topic changes. Therefore, for guided marker genes, it is also possible to probabilistically avoid updating *Z_m,n_* using a parameter.

#### Selection of genes for analysis

Consider a measured bulk gene expression matrix ***Y ∈ R^L×M^*** for *L* types of genes across *M* samples. In our proposed algorithm, collapsed Gibbs sampling is repeated *N* times, the sum frequency of *L* types of gene expression in all *M* samples. We excluded genes with that were outliers greater than 2σ from the log-normal distribution of gene expression levels. This procedure targets genes with consistently high expression levels observed in RNA-seq, notably mitochondrial and ribosomal genes. These genes are not informative and rather can introduce noise when modeling gene expression levels derived from changes in cell proportions. Furthermore, the transcriptome variation may also arise from simple expression change, such as those caused by perturbations. Therefore, in addition to the marker genes given as prior information, *topn* genes with large coefficient of variation among samples were selected for analysis. The hyperparameter *topn* was set to a default value of 100 for the analysis of blood samples and 1000 for more heterogeneous tissue samples. See Supplementary Notes S1 and *Hyperparameter sensitivity analysis* section for details.

#### Additional topics for tissue data analysis

Tissues are composed of a diverse range of cell populations. In addition to the guided topics that allow for cell identification, we also observe the effects of other topics, such as unexpected cell types or confounding between experiments. To account for these factors, we assume that *K* topics are composed of the *K_g_* guided topics of interest and the K_u_ additional unguided topics with no prior information.

To determine the optimal number of additional topics K_u_, we propose a recursive algorithm that tests gene contribution to each topic. After adding K_u_ unguided topics and performing deconvolution, we obtain *φ_k∈{1,2,…,k}_* ∈ *R^K×L’^*, which is the contribution of L′ types of non-marker genes to each additional topic. For each additional topic, we extract the top L′⁄K genes with highest contribution value and finally pool N_g_ genes. Then, we obtain Pearson correlation matrix *P ∈ R^Ku×K^* for the contribution profiles of the N_g_ genes across the added K_u_ topics and overall K topics. When an element of P is significantly positive, the added K_u_ topics are considered to contain redundancy. The value of K_u_ is increased until this redundancy is detected, and the final K_u_ is determined.

#### Strategy for robust estimation

The proportions of cell types in a sample are represented by the sample-topic distribution *θ_m_* (i = 1, ⋯ , *M*). Due to the dependence of the Gibbs sampling process on random variables, a single trial may not be sufficient for accurate estimation. To address this, we employed the median selection strategy, which integrates *θ*_m_ from multiple random seeds and ensures that the cell type proportions sum to 1.

#### Functional analysis for the additional topics

We can obtain K_u_ × En topic-gene distributions for unknown topics by assembling the output of the models considering K_u_ additional topics En times. K-means clustering was employed to determine the contribution of L genes to each potential additional topic. Subsequently, enrichment analyses of the top contributing L/K_u_ genes with Gene Ontology (GO) [24] and KEGG pathway [25] were performed utilizing the Fisher’s exact test. We also performed single-sample gene set enrichment analysis (ssGSEA) on the gene contribution values.

### Data preparation

To assess the performance of GLDADec in estimating gene expression levels in blood and tissue samples, we selected datasets that included corresponding bulk transcriptome and immune cell proportions determined by flow cytometry. Additionally, we obtained human clinical data for practical real-world data application. GLDADec takes a gene expression matrix ***Y ∈ R^L×M^*** for L genes across *M* samples as input and utilizes non-log linear scale data derived from the bulk transcriptome. A pseudo-bulk dataset derived from single-cell transcriptomes is also utilized to evaluate the robustness of GLDADec. A summary of the datasets used is provided in Supplementary Table S1. See Supplementary Notes S1 for details on the experimental conditions, including preprocessing of each dataset.

### Comparisons with other deconvolution methods

We compared GLDADec with seven competing bulk reference deconvolution methods, including FARDEEP, EPIC, CIBERSORT, DCQ, NNLS, RLR, and Elastic Net, utilizing their default settings where applicable [7–10]. Note that the input matrix for each method was normalized separately. Specifically, a log normalized gene expression matrix was used for DCQ and FARDEEP, while a non-log linear scaled matrix was utilized for the remaining methods. In addition, baseline scores for state-of-the-art methods including GTM-decon, BayesPrism, CIBERSORTx, MuSiC, and BSEQ-sc are reported by Swapna et al [14,20,26,27]. We obtained the deconvolution results from their GitHub repository (https://github.com/li-lab-mcgill/gtm-decon). See Supplementary Noste S2 for further details.

## Results

### Benchmarking with human blood samples

To begin with, we evaluated our algorithm using three datasets comprising bulk transcriptome data from human blood (GSE65133, GSE107572, GSE60424)[9,13,28,29], along with the corresponding proportions of immune cell types determined through flow cytometry. The marker gene names specific to each cell type were defined using domain knowledge, and 100 genes with large coefficients of variation (CV) among samples were included for analysis (as outlined in Supplementary File S1). Our proposed method, which utilizes marker gene names as prior information, achieved highly accurate predictions (Pearson correlations ranging from 0.39 to 0.99), considering each cell type individually (Figure 2A; Figures S1-3). Since the cell types with known proportions in these datasets are covered by LM22, a signature matrix defined by Newman et al [13], we compared the proposed GLDADec against seven bulk reference-based methods in using LM22 as a reference, including FARDEEP, EPIC, CIBERSORT, DCQ, NNLS, RLR, and ElasticNet [7–10], in terms of Pearson correlation and mean squared error (MSE). The proposed method demonstrated a high Pearson correlation across all benchmark datasets, and comparable or superior MSE scores to existing methods on datasets except for GSE60424 (Figure 2B; Figures S1-3,6A). Notably, for the dendritic cells of GSE107572, the estimation performance was significantly improved by our proposed method. The similarity matrix of the estimates across the three datasets revealed that GLDADec shows a relatively similar profile to DCQ (Figure S7A). The detailed experimental conditions for using existing methods are provided in Supplementary Note S2. To ensure that these estimates were robust and independent of the number of added high CV genes and hyperparameters, a sensitivity analysis was conducted (see *Hyperparameter sensitivity analysis* section).

**Fig. 2.**
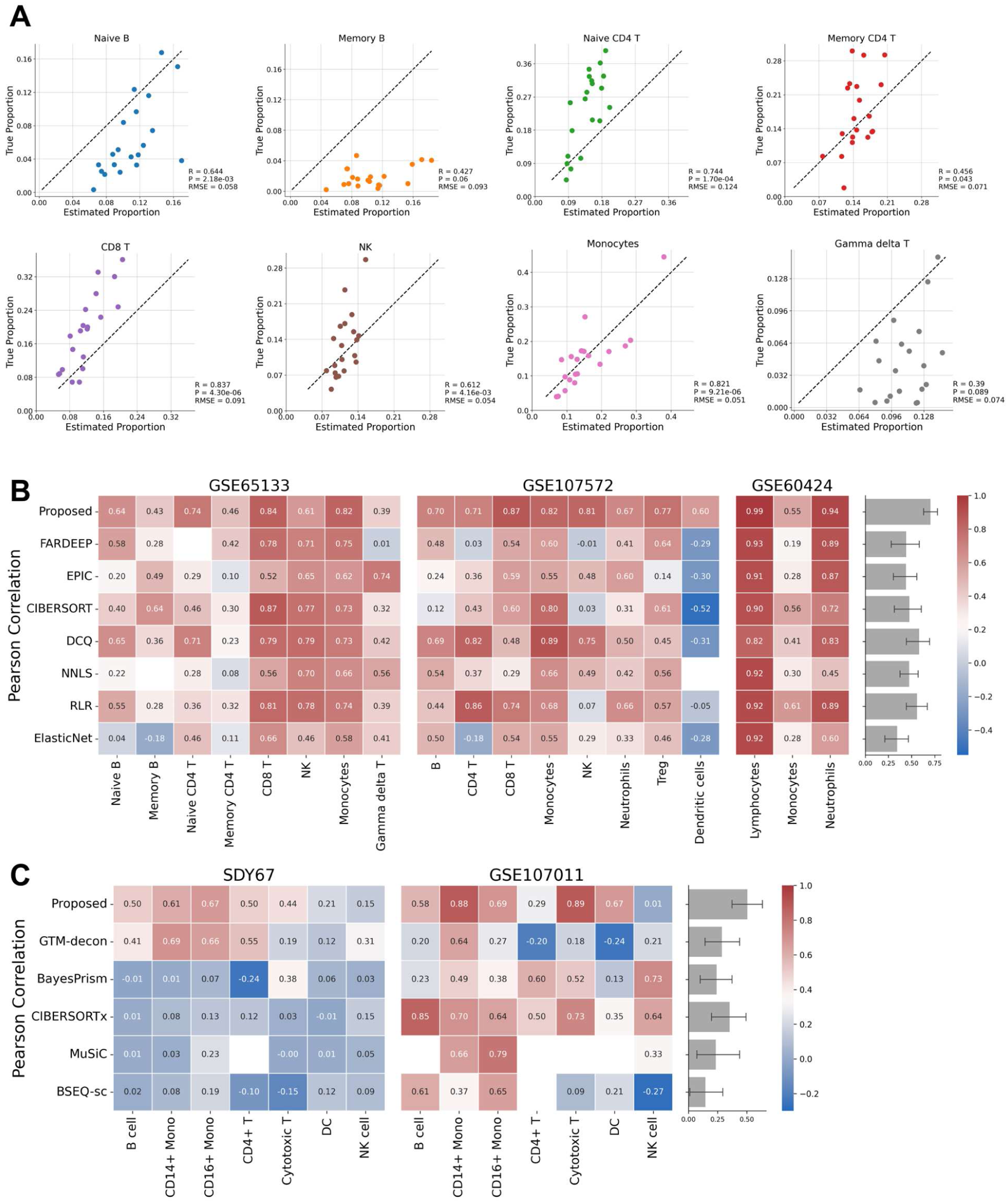
Benchmarking with human blood samples. **(A)** Scatterplots comparing estimated proportions and measured immune cell proportions in GSE65133. **(B)** Heatmaps showing Pearson correlation to compare the estimation performance against existing bulk reference-based methods using LM22 on three benchmark datasets, GSE65133, GSE107572, and GSE60424. **(C)** Heatmaps showing Pearson correlation to compare the proposed method to state-of-the-art methods on two benchmark datasets, SDY67 and GSE107011. The barplots on the right shows the performance of each method across all cell types.

We then conducted a benchmarking assessment of GLDADec against state-of-the-art methods, including GTM-decon, BayesPrism, CIBERSORTx, MuSiC, and BSEQ-sc [9,14,20,26,27]. These methods offer more detailed cell type estimations, utilizing single-cell RNA-Seq as prior information. SDY67 and GSE107011 [30,31], which maintain known detailed cell type proportions, were chosen as benchmark datasets, with baseline scores reported from Swapna et al [20]. While GTM-decon and CIBERSORTx exhibited relatively superior performance on SDY67 and GSE107011, respectively, GLDADec demonstrated equivalent or improved performance compared to existing methods across both datasets in terms of Pearson correlation and MSE (Figure 2C; Figure S4-5,6B). Analysis of the similarity matrix of the estimates revealed that GLDADec exhibits an independent profile compared to other methods (Figure S7B).

Taken together, these findings underscore the promise of semi-supervised topic modeling in deconvolution, with GLDADec enhancing estimation performance in real bulk scenarios derived from human blood.

### Impact of introducing additional topics

The transcriptome of tissues consists of heterogeneous cell populations, including cells that are not typically expected to be present in the tissue. To account for these unknown cell types, we performed modeling that treated them as additional topics. We investigated two scenarios to assess the utility of introducing additional topics into GLDADec.

Initially, we performed benchmarking on brain tissue data with known cell type proportions, known as ROSMAP [30]. All methods exhibited reasonably good performance on this dataset, with similar predictive capabilities. However, predicting endothelial cells and oligodendrocytes consistently posed challenges (Figure S8). This dataset served to assess whether a cell type excluded from guided topics could be reconstructed by introducing an additional topic. Remarkably, for all cell types except endothelial cells, the added topic proportions exhibited high correlation with the ground-truth proportions of the excluded cell type (Figure 3A). Furthermore, marker genes of the missing cell type were significantly enriched in the added topic as contributing genes (Figure 3B). To gain further biological insights into the additional topic, we conducted Gene Ontology (GO) [24] analysis for genes making substantial contributions to the added topic. Notably, the topic reconstructing microglia reflected the regulation of microglial differentiation and activation, while the topic reconstructing neurons reflected neurotransmitter transport as a biological process (Table 1; Table S2).

**Fig. 3.**
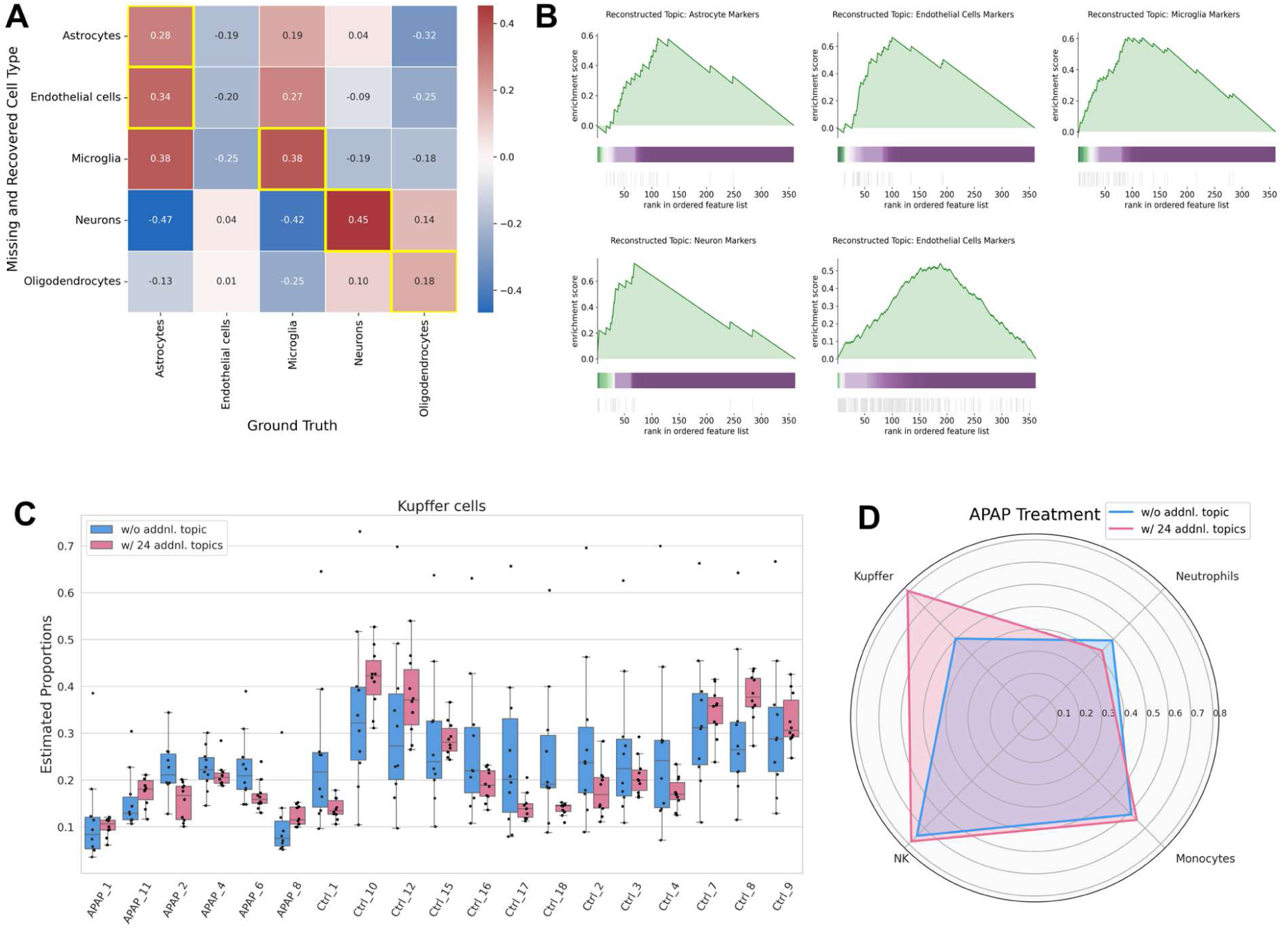
Assessment of the usefulness of incorporating additional topics in the analysis of tissue data. **(A)** Heatmap showing the correlation between the reconstructed cell type proportion and the ground-truth proportion. The maximum value for each row is highlighted in yellow. **(B)** Gene set enrichment analysis (GSEA) was performed on the gene list ranked by contribution to the additional topic reconstructing a missing cell type. The colored band represents the degree of contribution of each gene to the additional topic (green for a high contribution and purple for a low contribution). The bottom vertical black lines represent the location of the marker genes. **(C)** Variance of estimates at median selection for each sample with and without additional topics. **(D)** Radar chart comparing the difference in estimated performance with and without additional topics. The axis values signify the Pearson correlation between estimated and measured proportion of each immune cells.

**Table 1.**
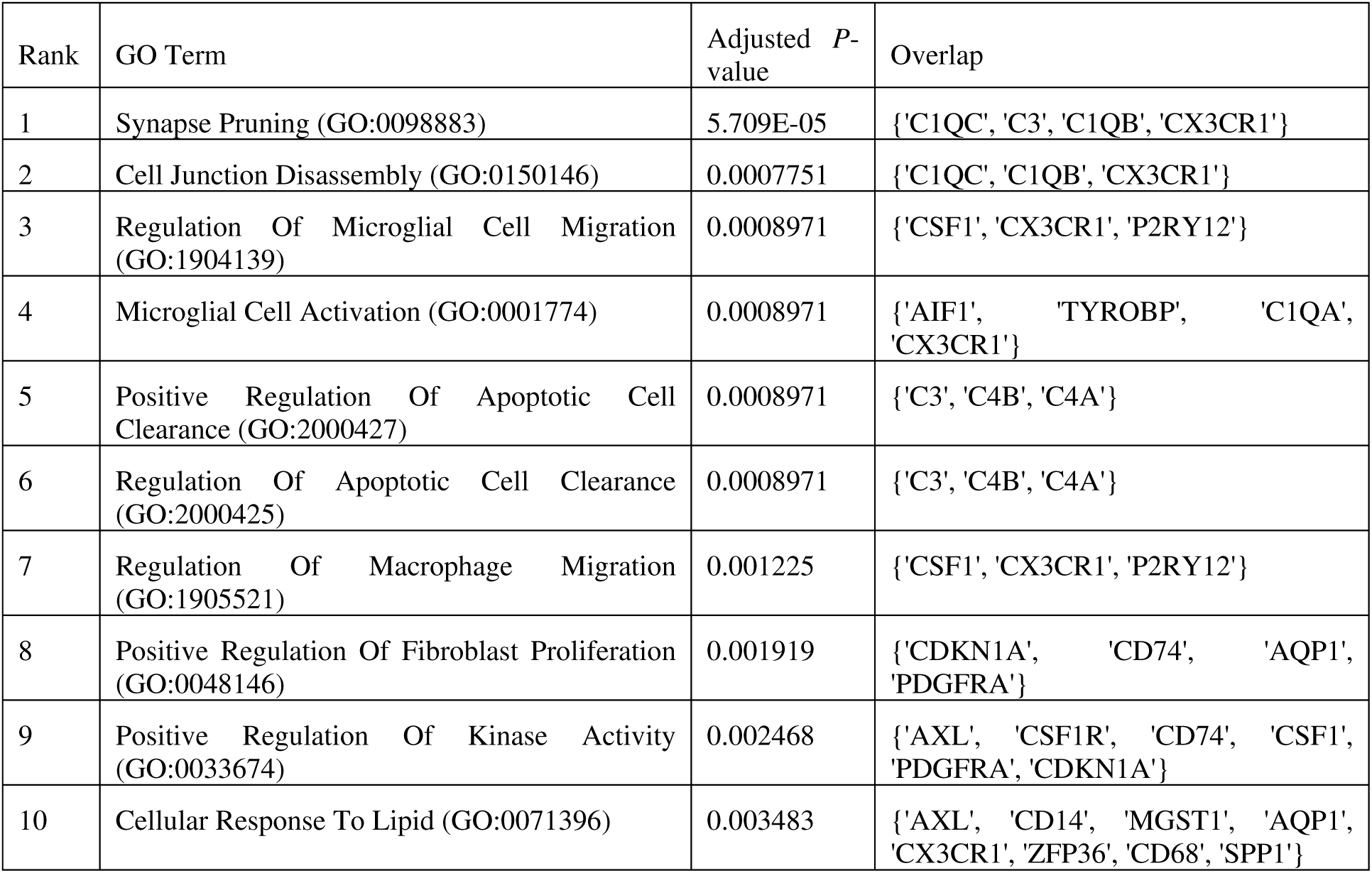
Gene Ontology (GO) enrichment analysis for the added topic that reconstructing missing microglia. The top 10 significantly enriched GO terms are shown with Benjamini–Hochberg adjusted *P*-values.

**Table 2.**
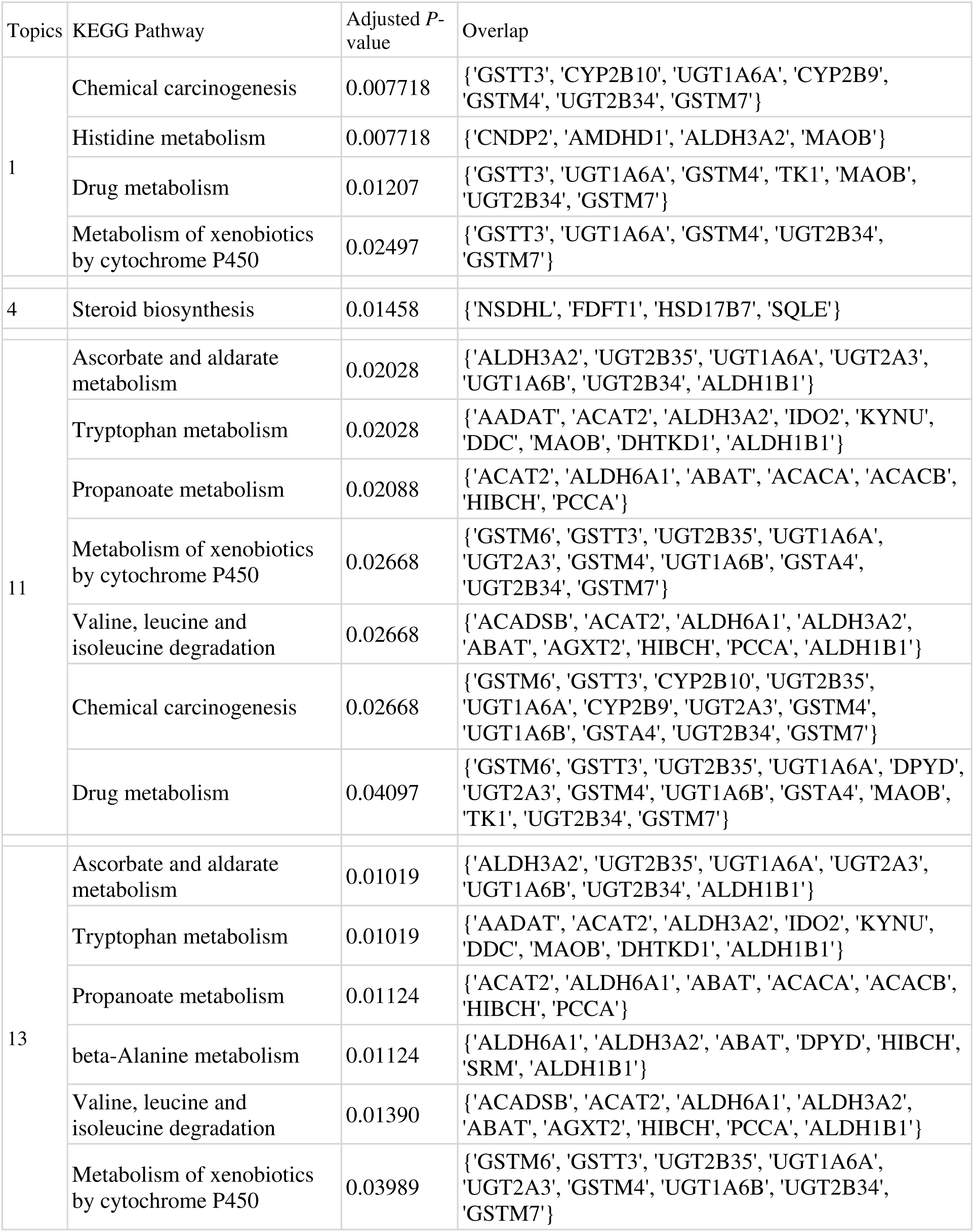
KEGG pathway enrichment analysis. Significantly enriched pathway names are shown with Benjamini–Hochberg adjusted *P*-values.

We next validated the impact of introducing additional topics by evaluating immune cell trafficking in tissues with perturbation, using liver bulk RNA-Seq data from various drug-induced liver injury mouse models. We compared modeling with and without additional topics, considering four known cell types (neutrophils, monocytes, NK cells, and Kupffer cells). The optimal number of additional topics increased with the number of genes analyzed for each acetaminophen (APAP) and alpha-naphthyl isothiocyanate (ANIT) treatment group (Figure S9A and B). Modeling with additional topics suppressed the variance of the estimates for several cell types, improving the performance of estimating the proportion of each cell type (Figure 3C; Figure S9C,10). In particular, the estimation of Kupffer cells with APAP administration was greatly improved (Figure 3D). To gain further biological insight into the additional topics, we conducted GO and KEGG pathway [25] analysis for the genes with large contribution to each additional topic. The significant biological processes were mainly related to metabolism and biosynthesis, consistent with a major biological role for liver tissue and reflecting the influence of potential transcriptomes such as hepatocytes (Figure S9D; Table 1; Table S2).

These results indicate that adding topics can improve the performance of guided cell estimation and allows the aggregation of biological functions that exist in the background of the target tissue as unknown topics.

### Comprehensive cell type analysis for mouse data

In recent years, databases containing marker genes have become prevalent and easily accessible [32,33]. By obtaining a data-driven approach to obtain marker gene names for a diverse array of cell types, we can accurately estimate the proportions of a comprehensive cell type.

In this study, we curated marker genes in mouse liver obtained from CellMarker [32] and defined marker genes specific to liver-related cell types (Supplementary File S3). Using these marker genes and applying the proposed method to liver tissue during drug-induced liver injury, we estimated the proportion changes for a wide range of 26 cell types (Figure 4A; Figure S11A). Notably, GLDADec allows the estimation of cell types such as hepatocytes and vascular smooth muscle cells (VSMCs), which have rarely been considered in conventional deconvolution methods. Pearson correlation was evaluated for four cell types validated by flow cytometry: neutrophils, monocytes, NK cells, and Kupffer cells. These cell types are widely recognized as playing a significant role in the pathophysiology of drug induced liver injury. As shown in Figures 4B and S11B, the proposed method accurately estimated the change in immune cell proportion with perturbation. In addition, although there were no ground-truths measured by flow cytometry for cell types such as hepatocytes and VSMCs, they showed significant positive or negative correlations with blood biochemistry values such as alanine aminotransferase (ALT) and aspartate aminotransferase (AST) (Tables S4, S5; Figure S12). These results suggest that accurate estimates that reflect individual responses can be achieved for a wide range of cell types.

**Fig. 4.**
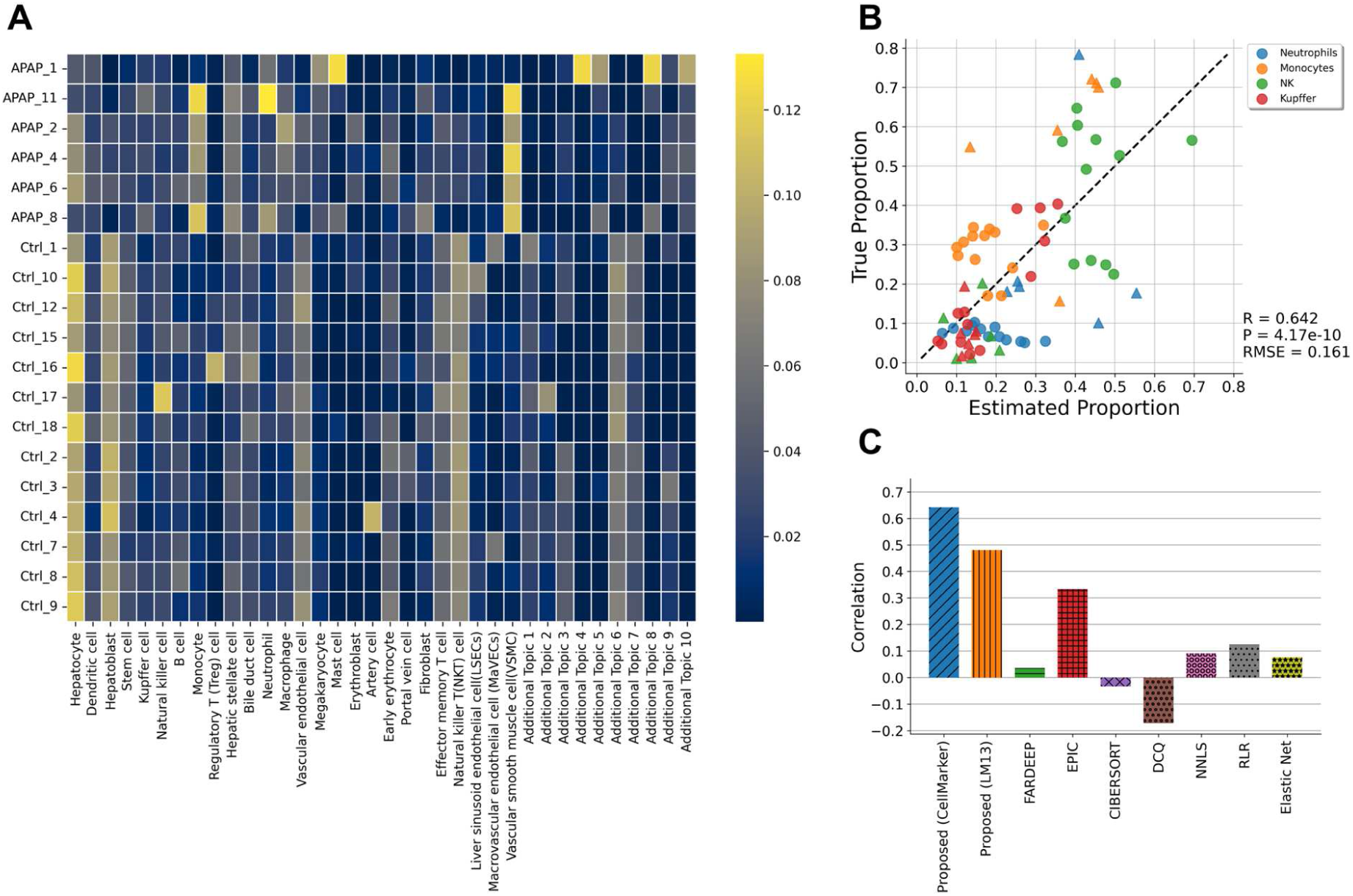
Prediction of immune cell trafficking in mouse liver tissue perturbed by acetaminophen (APAP). **(A)** Comprehensive cell type proportions estimation using marker gene names collected in data-driven manner. **(B)** Scatterplot showing the estimated proportions of immune cells vs. measured values in the same dataset. The circles indicate the control samples, and the triangles indicate the samples treated with APAP. **(C)** Comparison of performance using Pearson correlation with other notable deconvolution methods.

To assess the robustness of the proposed method to prior knowledge, we altered the marker genes used. Differentially expressed genes (DEGs) between LM13, 13 representative cell types of mouse liver, derived from transcriptome data of each cell type were deemed as marker genes and exhibited consistent superior estimation performance (Figure S13). Additionally, we evaluated the performance of existing reference-based methods utilizing DEGs, and the proposed method surpassed that of existing methods (illustrated in Figure 4C; Figure S11C).

### Application to rat data with poor marker information

Owing to the extensive use of rats in toxicology, there is plenty of toxicogenomic data for rats, a valuable resource for data analysis in drug and chemical safety assessments [17,34,35]. On the other hand, the cell marker information of rats was much lower than that of the mice. Our previous work showed that when using existing reference-based deconvolution, mouse-derived reference expressions are not extrapolatable, and rat-specific references should be used. However, databases containing gene expression profiles specific to each cell type in the rat are not as abundant as those in the mouse and must be obtained independently, which is costly and time-consuming.

In this section, we evaluate the performance of the proposed method using marker gene names defined in various scenarios. The method of defining the names of marker genes used to evaluate immune cell trafficking in the rat liver injury model was as follows:

1. Classical cell type markers for rats, mice, and humans reported by Natasha et al. [36]
2. Marker genes in the mouse liver obtained from CellMarker.
3. Names of DEGs derived from mouse LM6.

We found that the estimation performance was excellent when either was used as prior information (Figure 5A). Additionally, the estimation performance of the proposed method using each marker outperformed that of the existing reference-based method using mouse LM6, which is a set of representative immune cell types widely used in deconvolution (Figure 5B). These results suggest that the proposed method, which does not depend on the expression level of the reference gene, effectively achieves cell type proportion estimation using a priori information on the marker gene names that are conserved across species.

**Fig. 5.**
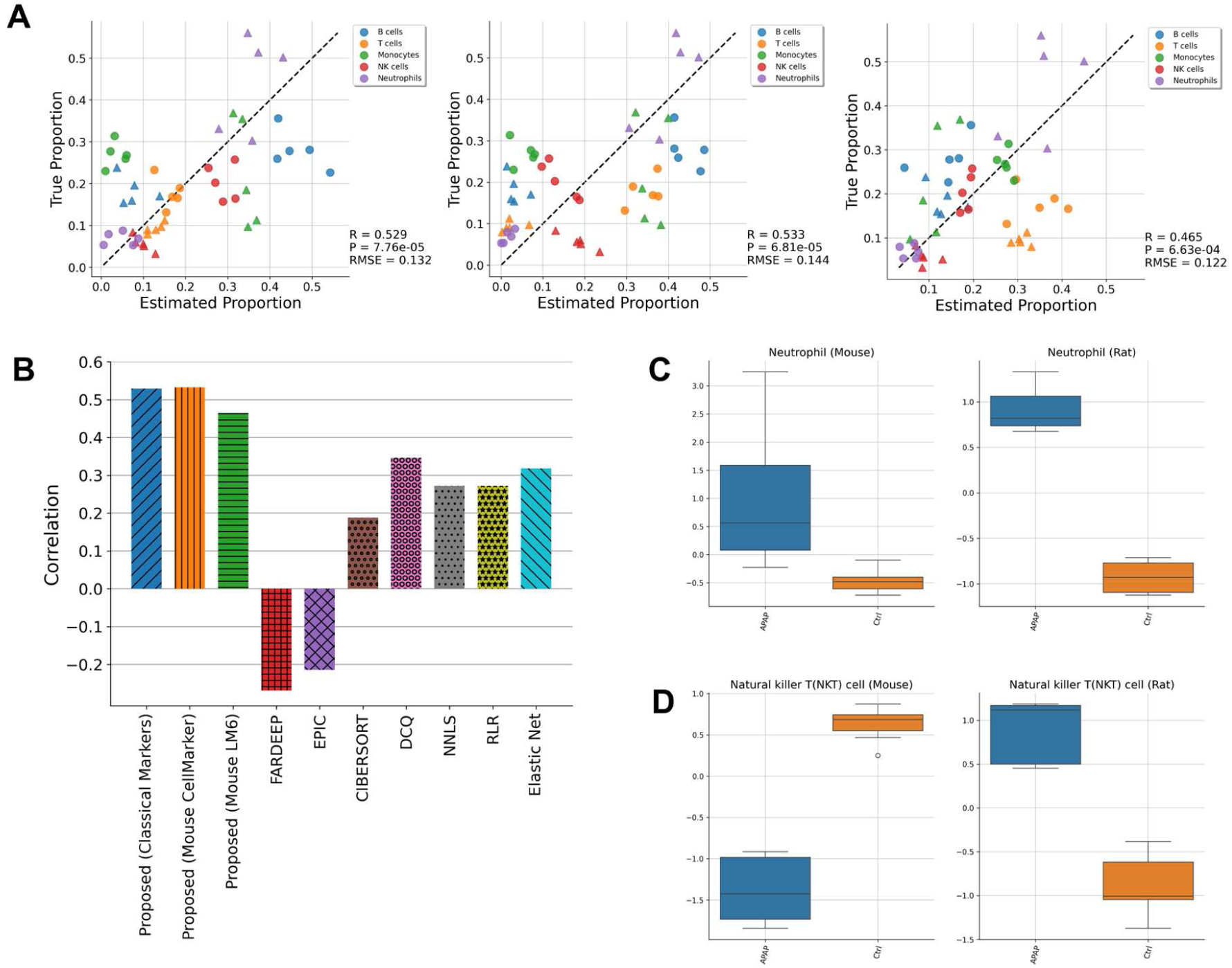
Estimation on rat samples with poor marker information. **(A)** Scatterplots showing the estimated proportions of immune cells alongside the corresponding measured values within the same dataset. These graphics represent the outcomes of defining marker genes in three distinct scenarios: classical marker genes, mouse marker genes obtained from CellMarker, and differentially expressed genes derived from cell type specific gene expression profiles in mice. The circular symbols denote control samples, while the triangular symbols represent samples that have been treated with acetaminophen (APAP). **(B)** Comparison of performance using Pearson correlation with other notable deconvolution methods. **(C-D)** Common and different patterns of immune cell trafficking in the liver between mice and rats treated with APAP.

Furthermore, our analysis of the immune response in rat liver injury using the GLDADec database provides a more extensive insight into the immune response in rat liver injury, utilizing the mouse marker gene names. By obtaining the same marker genes examined in mouse liver injury, we were able to estimate the comprehensive cell type proportions in rat liver injury and compare the profile changes between mice and rats (Figures S14A and B). Our findings revealed a striking similarity in the changes of cell proportions in mice and rats, such as an increase in neutrophils and monocytes and a decrease in hepatocytes, following the administration of acetaminophen (Figures 5C; Figure S14C). Additionally, we identified distinct immune cell trafficking patterns, including increased natural killer T (NKT) cells in mice but decreased in rats (Figure 5D). These conclusions indicate that our proposed method is a valuable tool for identifying similarities and differences in immune cell trafficking between species through comprehensive estimation of cell type proportions.

### Application GLDADec to tumor samples

The tumor tissue is composed of various cell types, including infiltrating immune cells, stromal and vascular cells, and subclonal cancer cells, as well as other cell types [37]. To validate our prediction in tumors, we applied GLDADec to 2037 tumor samples from The Cancer Genome Atlas (TCGA) for three tumor types: breast invasive carcinoma (BRCA), lung adenocarcinoma (LUAD), and liver hepatocellular carcinoma (LIHC) [38–41]. We used marker gene names obtained from CellMarker as prior information for each cancer subtype in the background tissues. Our proposed method stratified the cancer subtypes by comprehensively estimating the proportion of various cell types, including tissue-specific cells such as ciliated cells in LUAD samples and hepatocytes in LIHC samples (Figure 6A).

**Fig. 6.**
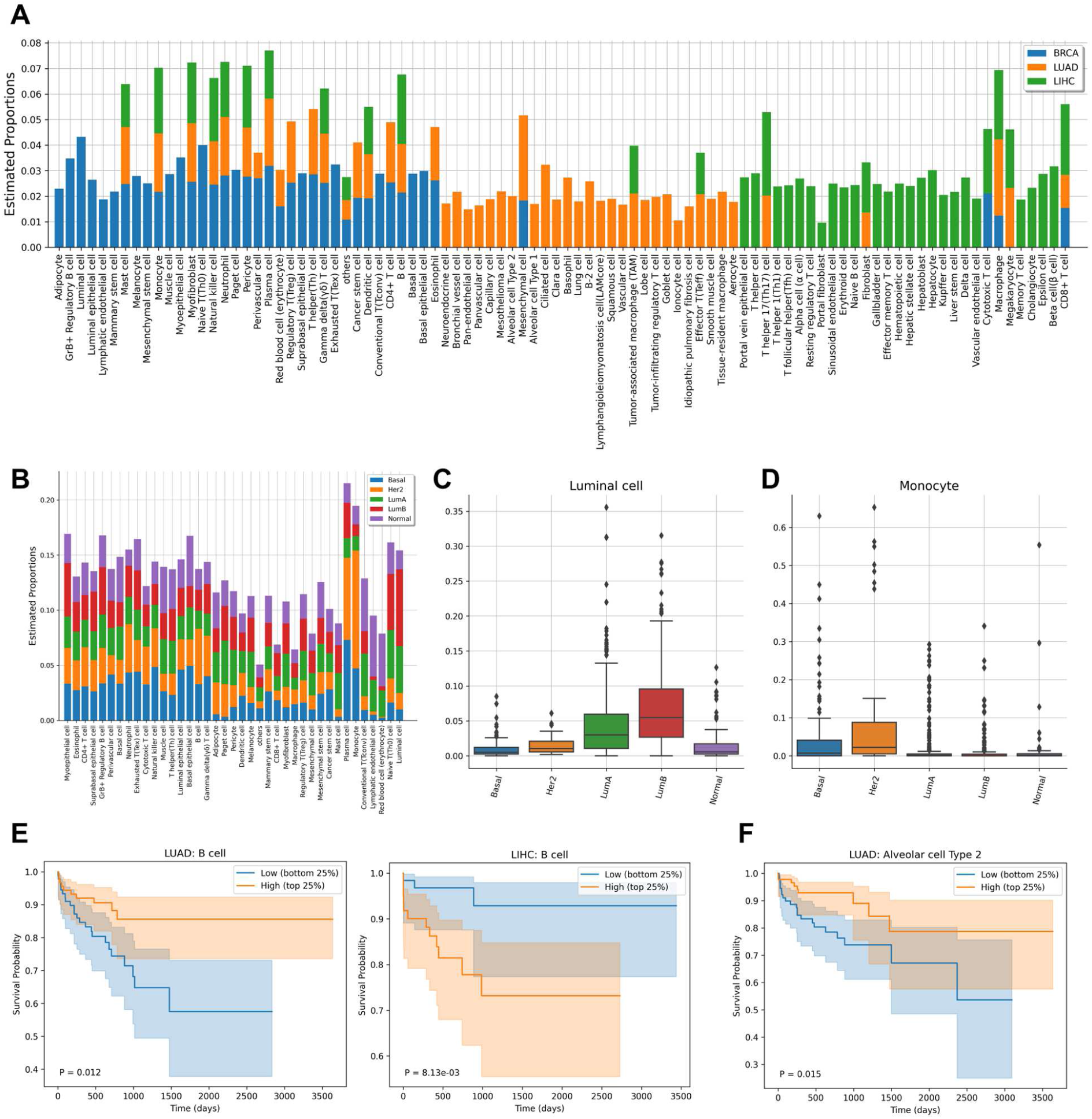
Application of GLDADec to tumor samples. **(A)** Stacked barplots showing the estimated comprehensive cell type proportions in each tumor samples derived from BRCA, LUAD, and LIHC. **(B)** Relative proportions of comprehensive cell types in different subtypes of BRCA. **(C-D)** Boxplots showing specific accumulation and infiltration between each subtype of BRCA. **(E-F)** Kaplan-Meier plots displaying the relationship between survival and the infiltration of B cells in LUAD and LIHC, as well as Alveolar cell type 2 in LUAD. The patients were divided into two groups based on the top and bottom 25^th^ percentiles of the target cells, and the results were analyzed using the log-rank test. The transparent colors in the plots represent 95% confidence bands, and the *P*-values were computed using the log-rank test.

To assess the refined stratification performance more comprehensively, we examined the differences in estimated cell proportions for each breast cancer subtype, as reported by Brian et al. [42] (Figure 6B). As anticipated, the proportion of luminal cells was found to be higher in subtypes LumA and LumB (Figure 6C). Moreover, we observed a distinct pattern of immune cell infiltration, particularly monocytes, B cells, and plasma cells, in the Her2 subtype (Figure 6D; Figure S15). This suggests that subtype classification based on cell proportions estimated by GLDADec is feasible and may offer novel insights into the unique migration patterns of immune cells within each subtype.

Next, we examined the relationship between the predicted proportion of cell types and patient survival. Tumor samples were categorized according to the estimated proportion of specific cell types, and Cox proportional hazards regression was employed to determine survival rates between the top and bottom 25% of samples. Our analyses revealed that patients with lung adenocarcinoma (LUAD) exhibiting a high level of B cell infiltration had worse overall survival (*P* = 0.012), while those with liver hepatocellular carcinoma (LIHC) demonstrated improved survival (*P* = 8.13e-03) (Figure 6E). These findings indicate that B cells contribute differently to the prognosis and treatment of breast and liver cancers, which is consistent with previous reports [43,44]. Moreover, our study found that the infiltration of immune cells, such as γδT and CD4+ T cells, was associated with better clinical outcomes in breast cancer, which supports previous studies [45,46] (Figure S16). In addition, GLDADec can estimate the proportion of various cell types with marker gene names, and it is expected to provide novel insights. For instance, we reported that the loss of alveolar type II cells may be the cause of the poor prognosis of LUAD (Figure 6F).

### Hyperparameter sensitivity analysis

The benchmark dataset was utilized to carry out sensitivity analyses for the hyperparameters *α* for sample-topic distribution, *η* for topic-gene distribution, as well as the number of high CV genes and the number of median selection attempts. It was observed that GLDADec exhibits robustness with respect to *α* and *η* under conditions between 0 and 1, which are commonly used in LDA [21,47]. Under conditions that allow for more than one, the estimates were found to be less dependent on *α* and more susceptible to changes in *η* (Figure S17A). In terms of the number of high CV genes to add, it was observed that gamma-delta T cells (γδT) exhibited better estimation performance when a higher number of genes were added; however, this effect was limited for many other cell types (Figure S17B). Regarding the number of median selection attempts, there was little effect on the human blood-derived benchmark dataset, while more accurate estimation performance was obtained by increasing the number of trials in the analysis on perturbed mouse and rat tissues (Figure S17C and D).

### Evaluation of robustness using pseudo bulk dataset

Given that topic modeling for a limited number of documents remains a challenging task [48,49], we examined the influence of sample size on the estimation performance of GLDADec. To simulate tissue data for varying sample sizes, pseudo lung bulk data was generated from single-cell RNA-Seq data. GLDADec was applied to 100 randomly generated samples, demonstrating its capability to yield highly accurate deconvolution predictions for both immune cells and tissue background cells (Figure S18A). When the number of samples was reduced to 10 or 5, estimation performance slightly decreased, and variance increased for certain cell types, such as alveolar type I cells and cytotoxic T cells. Nonetheless, the overall estimation performance remained high across immune cell subtypes (Figure S18B).

We further verified the robustness of GLDADec by ensuring its independence from differences in the distribution of expression levels between the reference and the bulk samples under analysis. This was achieved by partitioning the dataset used for pseudo bulk creation into main and test sets without any information leaks, and utilizing the reference defined from the test cells. Consequently, high estimation performance was attained, even with conventional methods such as ElasticNet and RLR (Figure S19A). Conversely, when lung single-cell RNA-Seq data from a distinct dataset was utilized as a reference [50], these reference-based methods consistently exhibited inferior performance (Figure S19B).

Overall, we demonstrated the robustness of our proposed method to sample size and prior information through analyses conducted on pseudo bulk data.

## Discussion

We introduce GLDADec, a marker gene name-based deconvolution approach, to estimate the cell type proportions present in a heterogeneous sample. In the process of inferring the topic distribution of each sample, we utilize marker gene names specific to each cell type as partial prior information to guide the cell name to the topic. Additionally, we can detect and incorporate unknown potential topics that characterize the sample, alongside the guided topics, simultaneously. The estimated topic distribution can be utilized to calculate the proportion of cell types from the bulk RNA-Seq dataset.

A notable advantage of GLDADec is the ease of accessing the name of the marker gene used as prior information. Although gene expression profiles of immune cells can typically be obtained in humans and mice, this is not always feasible for species such as rats that are used in limited areas or for tissue-specific parenchymal cells [35]. As a result, reference-based methods are dependent on the amount of prior information available for the scope of their analysis. However, marker gene names are uniformly accessible from databases, and there are also classical markers that are conserved across species, enabling a far more extensive range of prior information to be utilized in terms of the number of cell types and species to be analyzed.

One of the benefits of this method is its ability to infer additional topics other than guided topics, which may be useful in accounting for the impact of unknown cell types or removing confounding factors such as batch effects. In our analysis of liver samples from a drug-induced liver injury mouse model, we observed that the estimation of Kupffer cells was significantly improved by considering additional topics, whereas the benefits for neutrophils were minimal (Figure 3D). For neutrophils, the estimated values were robust even without considering additional topics, suggesting that the variance in neutrophils was small and thus the impact of additional topics on the cell type was limited (Figure S10A). We also found that the additional topics reflected biological characteristics, such as enrichment in metabolic and biosynthetic pathways (Figure S9D), which is consistent with the biological role of the liver. These topics likely reflect the influence of the transcriptome derived from hepatocytes, which constitute the majority of liver tissue.

Applying GLDADec, we quantified the fraction of diverse cell types within tumor samples registered in the TCGA database. We focused on minor cell types that were not typically identified through conventional approaches. For instance, we revealed the significant association between the presence of alveolar type II cells and unfavorable clinical outcomes for patients with LUAD (Figure 6F). These findings possess the potential to facilitate subsequent analyses and contribute novel perspectives.

It should be noted that the cell types covered by GLDADec are contingent upon the extent of knowledge of cell types and their marker genes that have been accumulated in the field of life sciences. In this study, we obtained marker gene names from domain knowledge or CellMarker database. However, the accuracy of marker gene selection may decrease when the labeling of the ground-truth is imprecise, as seen in the case of “lymphocytes” in GSE60424. This may explain why GLDADec exhibited a poor MSE score in GSE60424. Furthermore, although the CellMarker database is well characterized, there is room for improvement in coverage, as eosinophils, a cell type that plays a protective role in acute liver injury, are not registered in liver tissue [51]. Therefore, for a more comprehensive analysis, it may be necessary to integrate multiple databases that store marker genes to consolidate information. On the other hand, it’s worth noting that the protein markers listed in these databases may not consistently serve as good transcriptional markers. Currently, data cleansing relies heavily on manual curation employing domain knowledge. Given the recent advancements in large language models, extracting marker gene information from scientific literature could prove beneficial in identifying appropriate marker gene names for deconvolution of tissues of interest. While social acceptance is crucial as knowledge, such knowledge miners could prove to be powerful partners of the proposed method.

## Key Points

- We introduce GLDADec, a novel bulk deconvolution method leveraging marker gene names as partial prior information for estimating cell type proportions.
- GLDADec adopts a semi-supervised learning algorithm, utilizing cell type marker genes to address challenges present in both conventional reference-based and reference-free methods simultaneously.
- GLDADec demonstrates strong estimation performance and biological interpretability as evidenced by benchmarking across blood-derived and perturbed tissue datasets.
- We utilized GLDADec on TCGA tumor samples, conducting cancer subtype stratification and survival analysis, showcasing its utility in clinical data analysis.

## Declarations

### Ethics approval and consent to participate

The studies reported in this article were performed in accordance with the guidelines provided by the Institutional Animal Care Committee (Graduate School of Pharmaceutical Sciences, the University of Tokyo, Tokyo, Japan).

### Consent for publication

Not applicable.

### Availability of data and materials

Code, models, and data are available at https://github.com/mizuno-group/GLDADec.

### Contributions

IA: Conceptualization, Data curation, Formal analysis, Methodology, Software, Investigation, Writing – Original draft, Visualization.

TM: Conceptualization, Resources, Supervision, Project administration, Writing – Original draft, Writing – Review and editing, Funding acquisition.

HK: Writing – Review and editing.

## Algorithms

**Algorithms 1:**
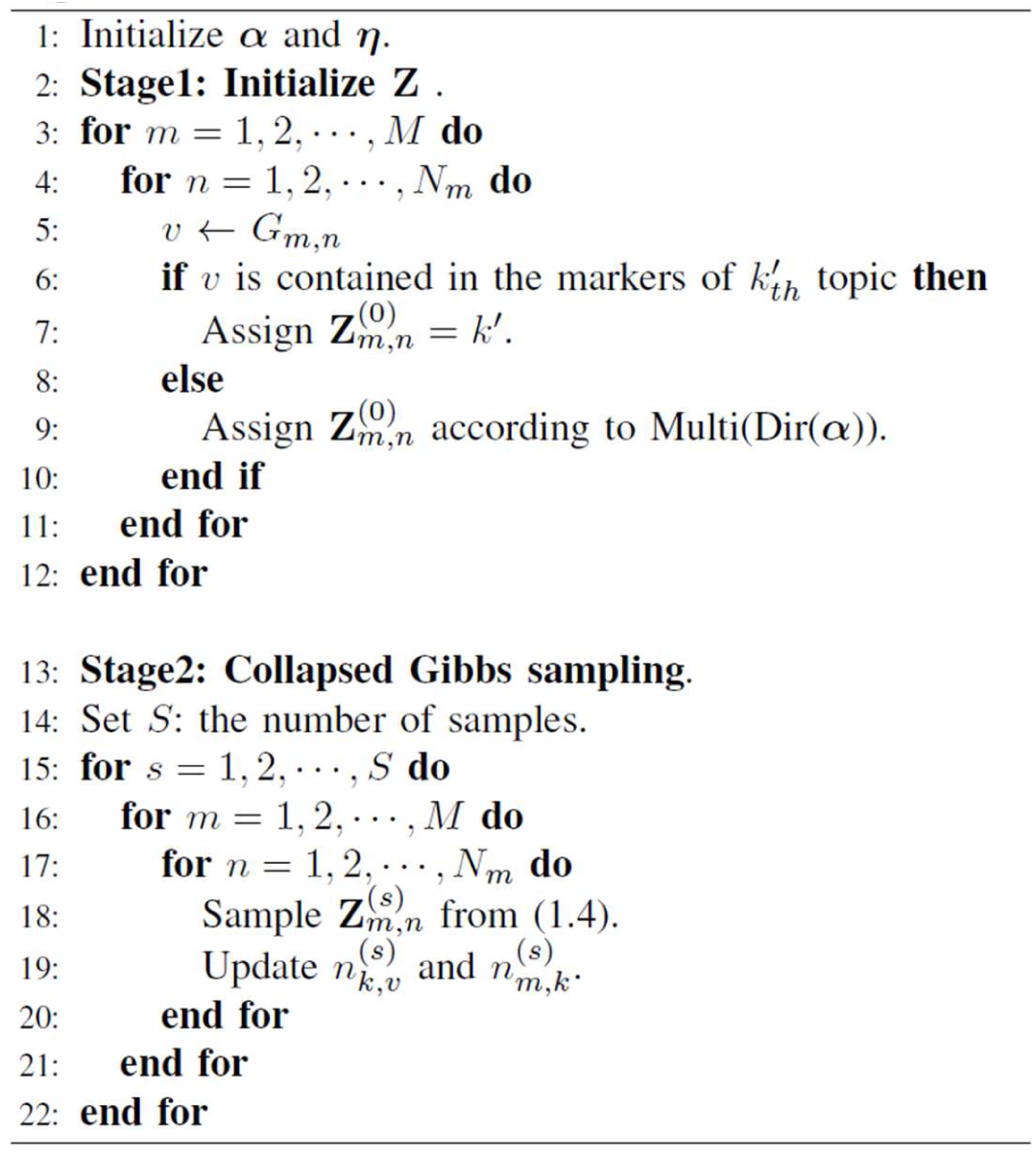
Overall Inference Process of GLDADec

**Figure S1.**
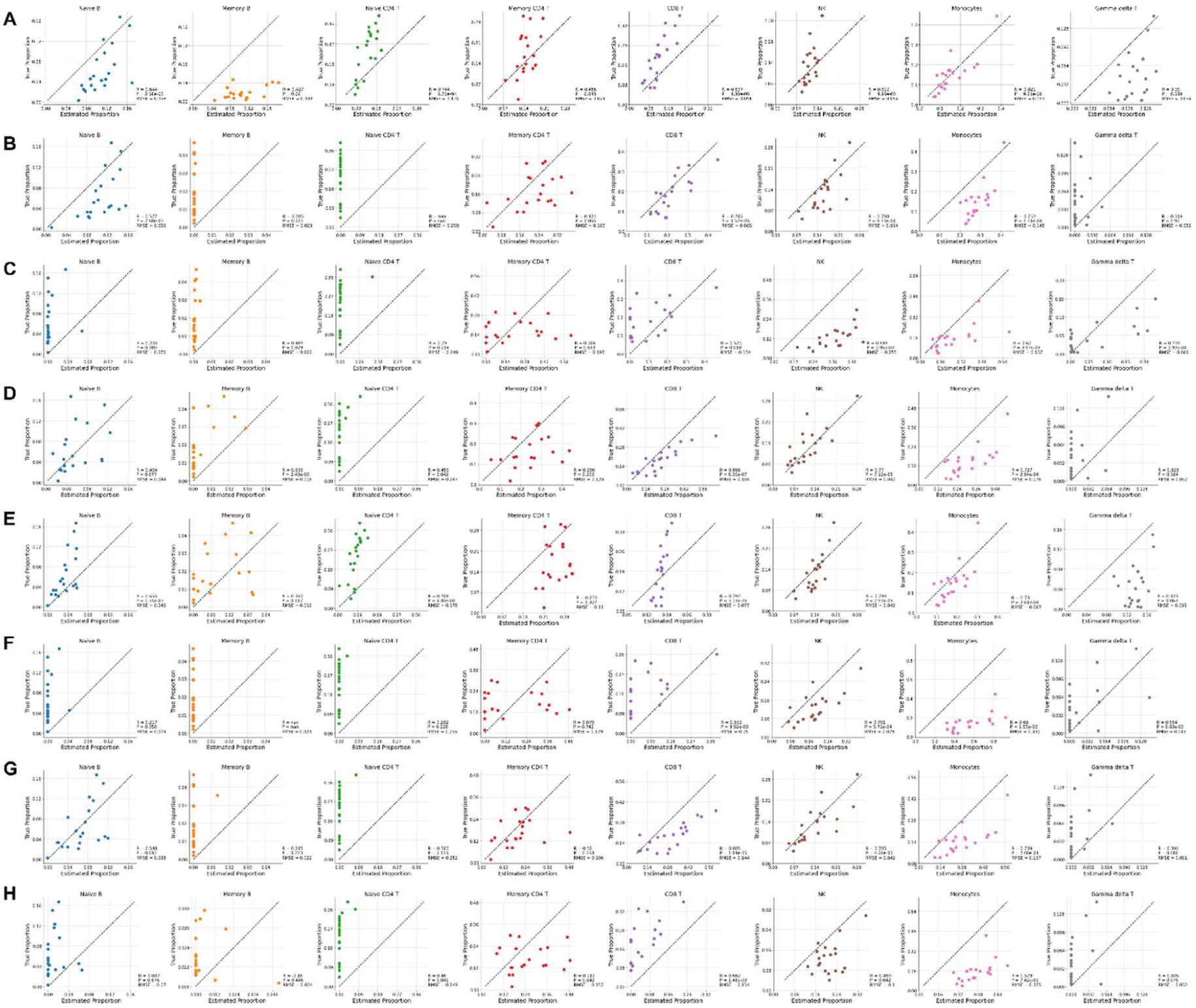
Scatterplots showing the correlation between estimated proportion and ground-truth proportion in the benchmark dataset GSE65133. Each row represents deconvolution method, (A) GLDADec, (B) FARDEEP, (C) EPIC, (D) CIBERSORT, (E) DCQ, (F) NNLS, (G) RLR, and (H) ElasticNet, respectively.

**Figure S2.**
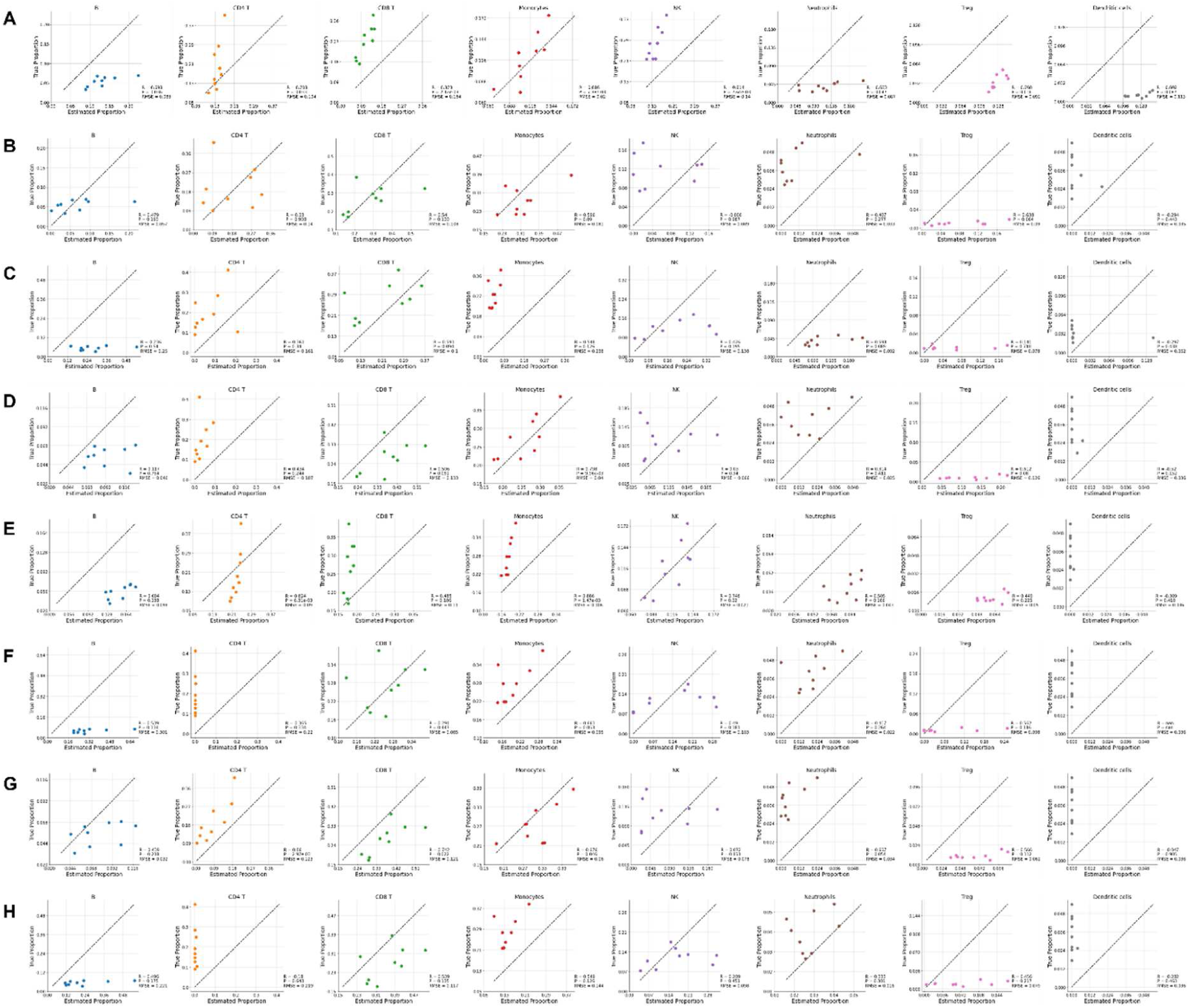
Scatterplots showing the correlation between estimated proportion and ground-truth proportion in the benchmark dataset GSE107572. Each row represents deconvolution method, (A) GLDADec, (B) FARDEEP, (C) EPIC, (D) CIBERSORT, (E) DCQ, (F) NNLS, (G) RLR, and (H) ElasticNet, respectively.

**Figure S3.**
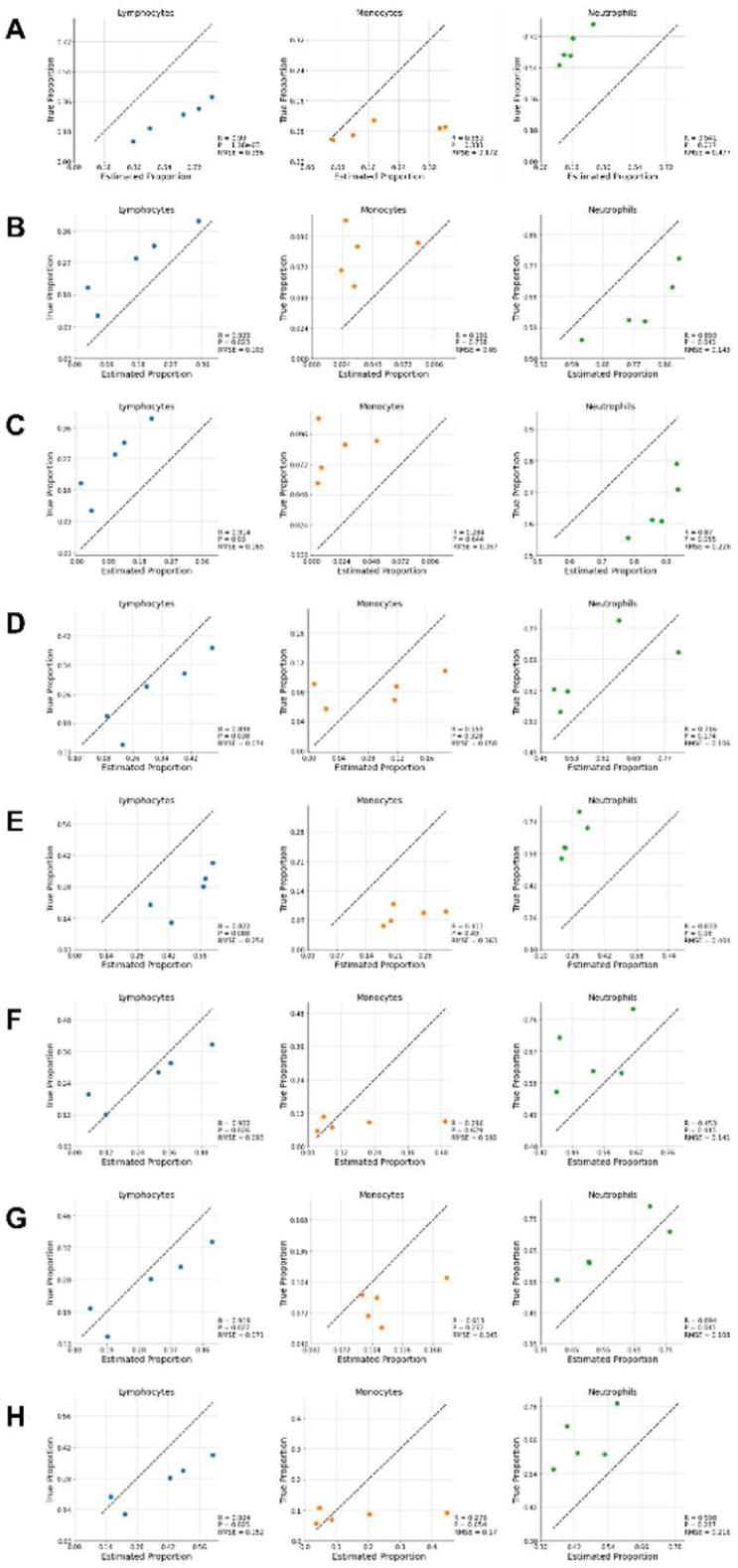
Scatterplots showing the correlation between estimated proportion and ground-truth proportion in the benchmark dataset GSE60424. Each row represents deconvolution method, (A) GLDADec, (B) FARDEEP, (C) EPIC, (D) CIBERSORT, (E) DCQ, (F) NNLS, (G) RLR, and (H) ElasticNet, respectively.

**Figure S4.**
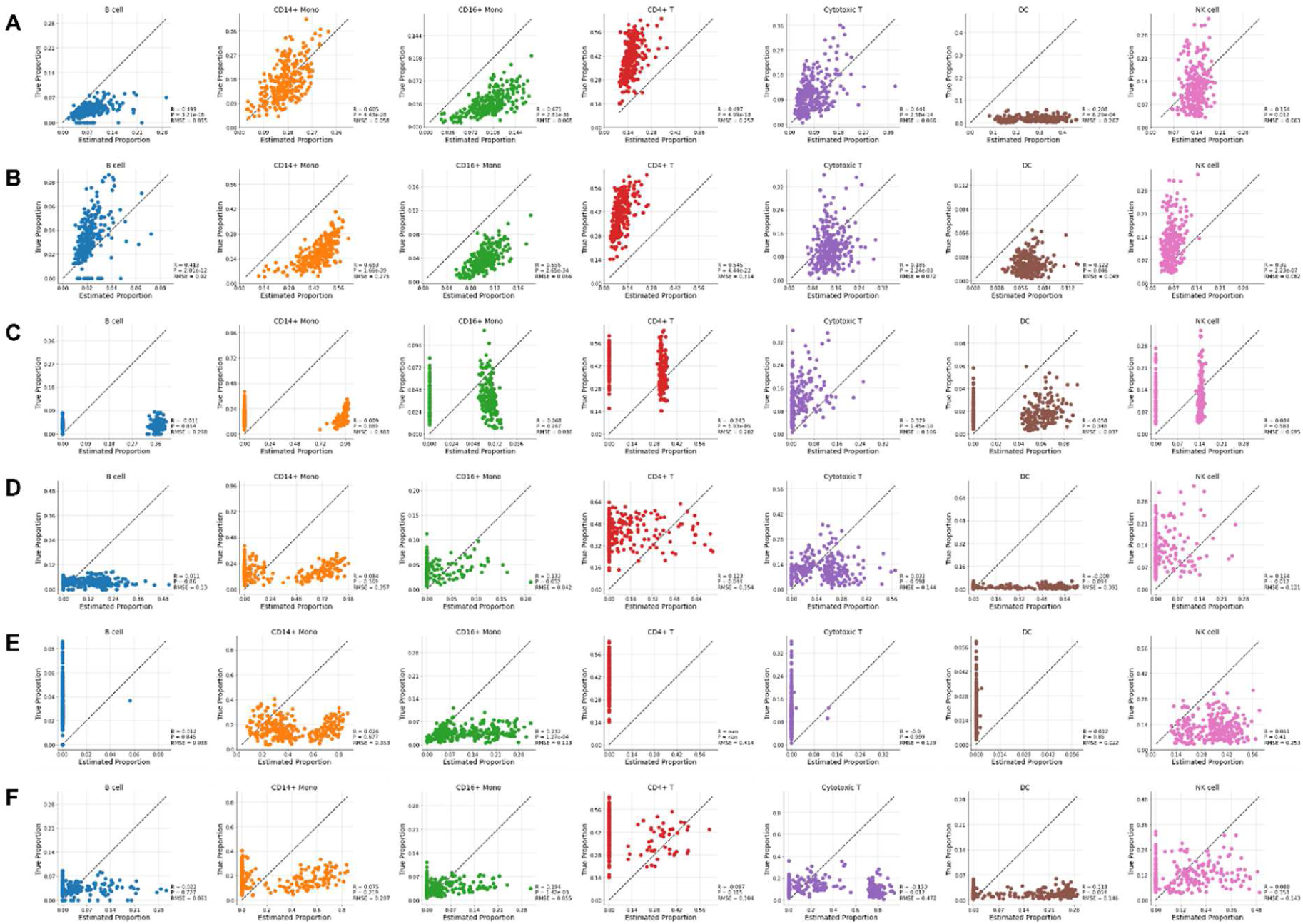
Scatterplots showing the correlation between estimated proportion and ground-truth proportion in the benchmark dataset SDY67. Each row represents deconvolution method, (A) GLDADec, (B) GTM-decon, (C) BayesPrism, (D) CIBERSORTx, (E) MuSiC, and (F) BSEQ-sc, respectively.

**Figure S5.**
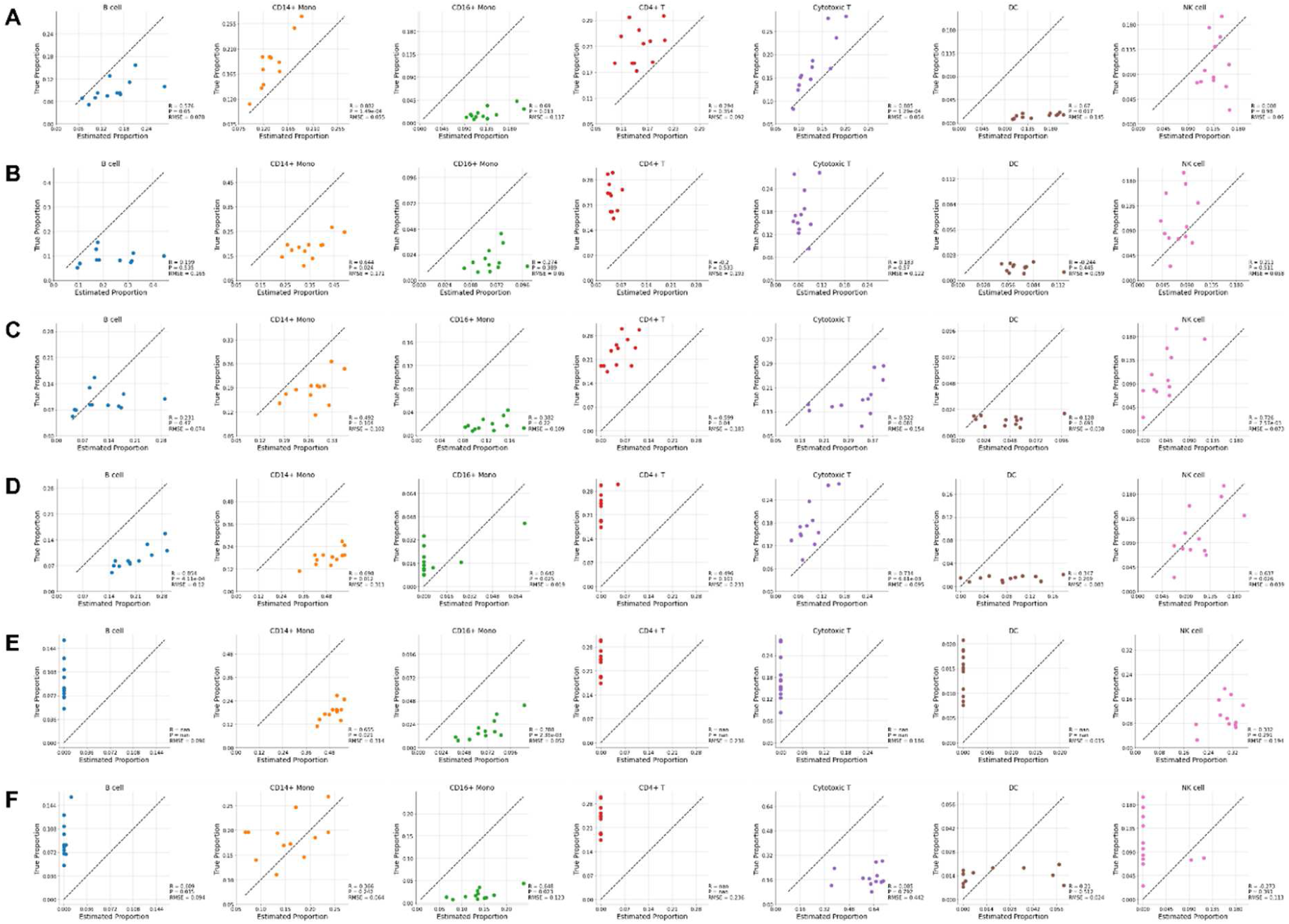
Scatterplots showing the correlation between estimated proportion and ground-truth proportion in the benchmark dataset GSE107011. Each row represents deconvolution method, (A) GLDADec, (B) GTM-decon, (C) BayesPrism, (D) CIBERSORTx, (E) MuSiC, and (F) BSEQ-sc, respectively.

**Figure S6.**
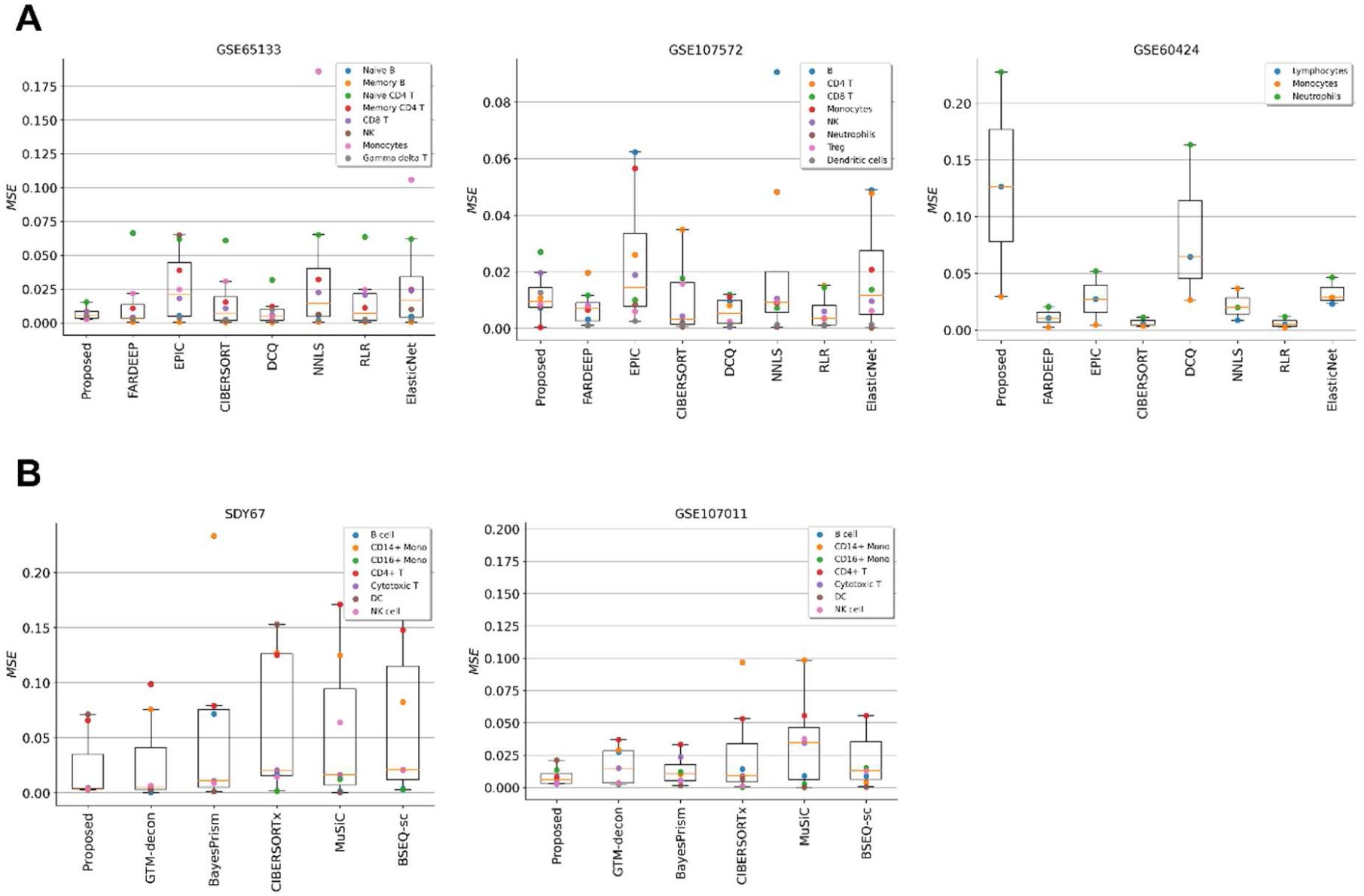
(A) Boxplots showing the mean square error (MSE) to compare the estimation performance against existing bulk reference-based methods using LM22 on three benchmark datasets, GSE65133, GSE107572, and GSE60424. (B) Boxplots showing the mean square error (MSE) to compare the estimation performance against the proposed method to state-of-the-art methods on two benchmark datasets, SDY67 and GSE107011. Each box extends from the 25^th^ percentile (bottom) to 75^th^ percentile (top), and the whisker indicates the farthest data point within 1.5-fold of inter-quartile range.

**Figure S7.**
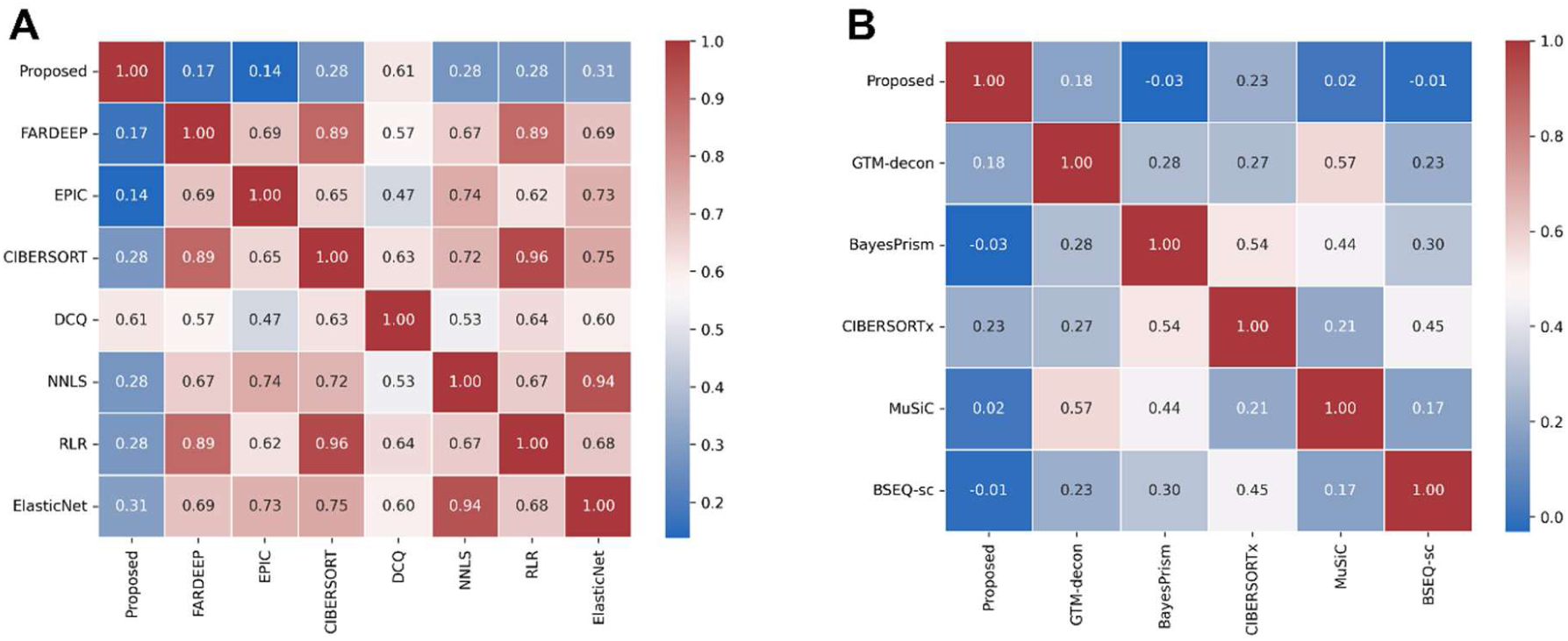
Similarity matrix of estimated values among (A) traditional bulk reference-based methods and (B) state-of-the-art methods.

**Figure S8.**
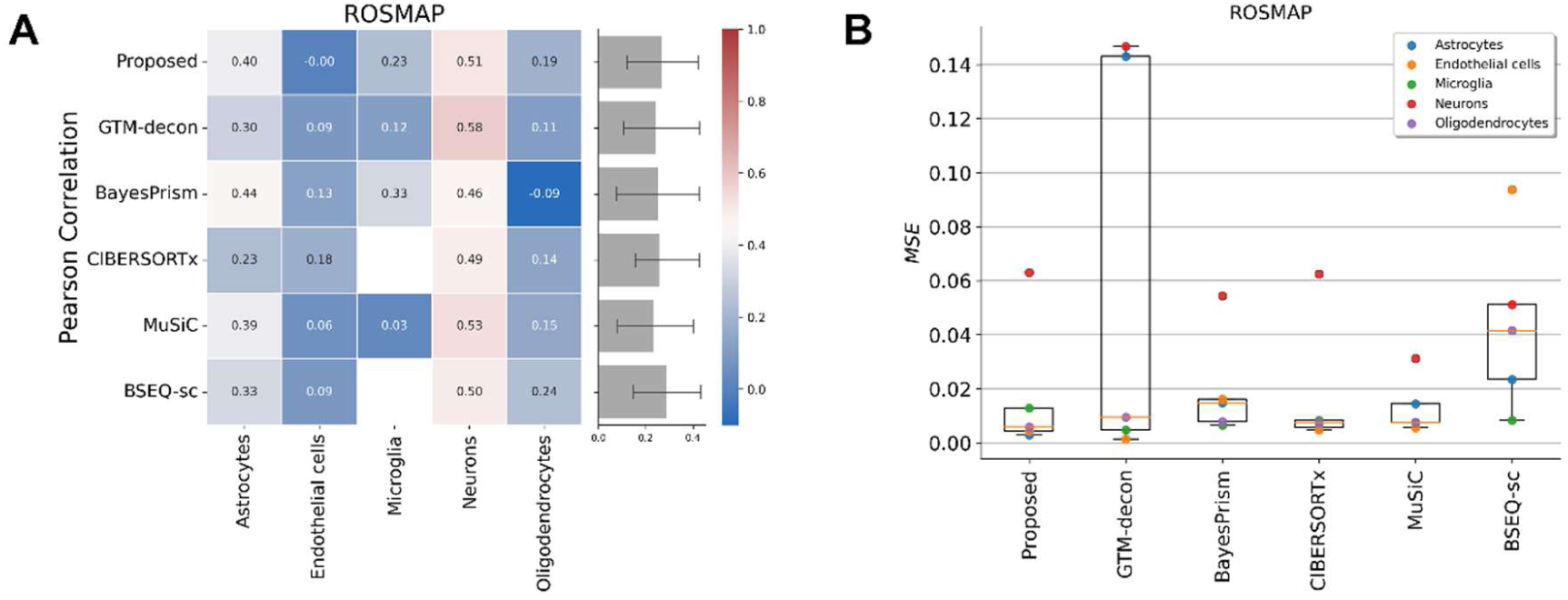
Benchmarking with human brain prefrontal cortex bulk dataset. (A) Heatmaps of comparing the estimation performance against state-of-the-art methods for Pearson correlation. The barplots on the right shows the performance of each method across all cell types. (B) Boxplots showing the mean square error (MSE) to compare the estimation performance against the proposed method to state-of-the-art methods. Each box extends from the 25^th^ percentile (bottom) to 75^th^ percentile (top), and the whisker indicates the farthest data point within 1.5-fold of inter-quartile range.

**Figure S9.**
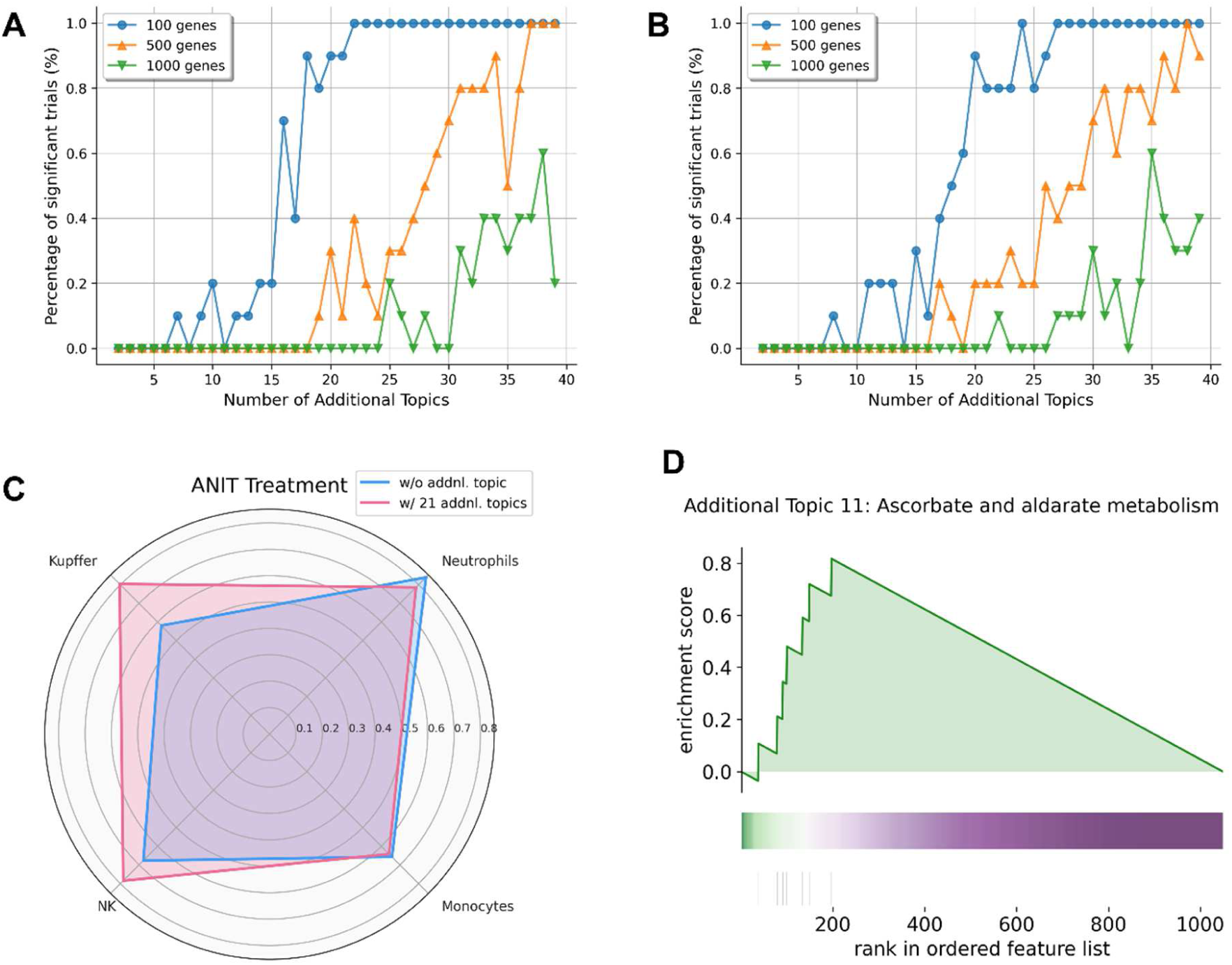
Evaluation of the usefulness of additional topics in the analysis of perturbed tissue data. The relationship between the number of additional topics that are significantly independent of each other and the number of genes to be analyzed for (A) acetaminophen (APAP)-perturbed and (B) alpha-naphthyl isothiocyanate (ANIT)-perturbed liver tissue. (C) A Radar chart comparing the difference in estimated performance with and without the additional topics for ANIT-perturbed liver tissue. The axis values indicate the Pearson correlation between estimated and measured proportion of each immune cells. (D) Gene set enrichment analysis (GSEA) was carried out on the ranked list of genes ranked according to their contribution to Topic 11. The colored band signifies the extent of each gene’s contribution to the additional topic 11, with green representing a high contribution and purple indicating a low contribution. The bottom vertical black lines indicate the position of the genes that overlap with the pathway.

**Figure S10.**
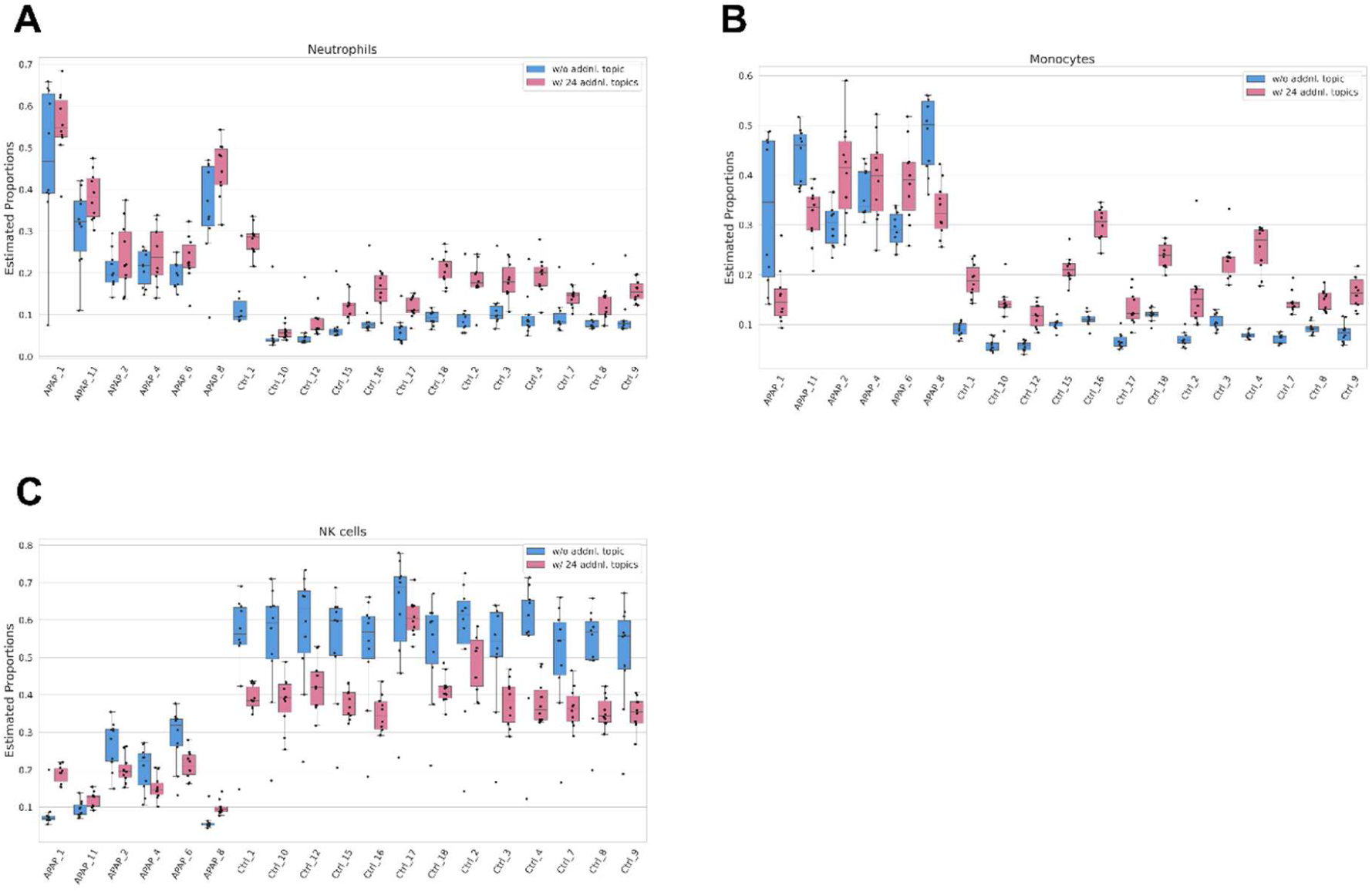
Variance of the estimate at ensemble in each sample with and without additional topics considered. Bar plots showing the estimated values of **(A)** neutrophils, **(B)** monocytes, and **(C**) Natural killer cells in each sample after APAP administration.

**Figure S11.**
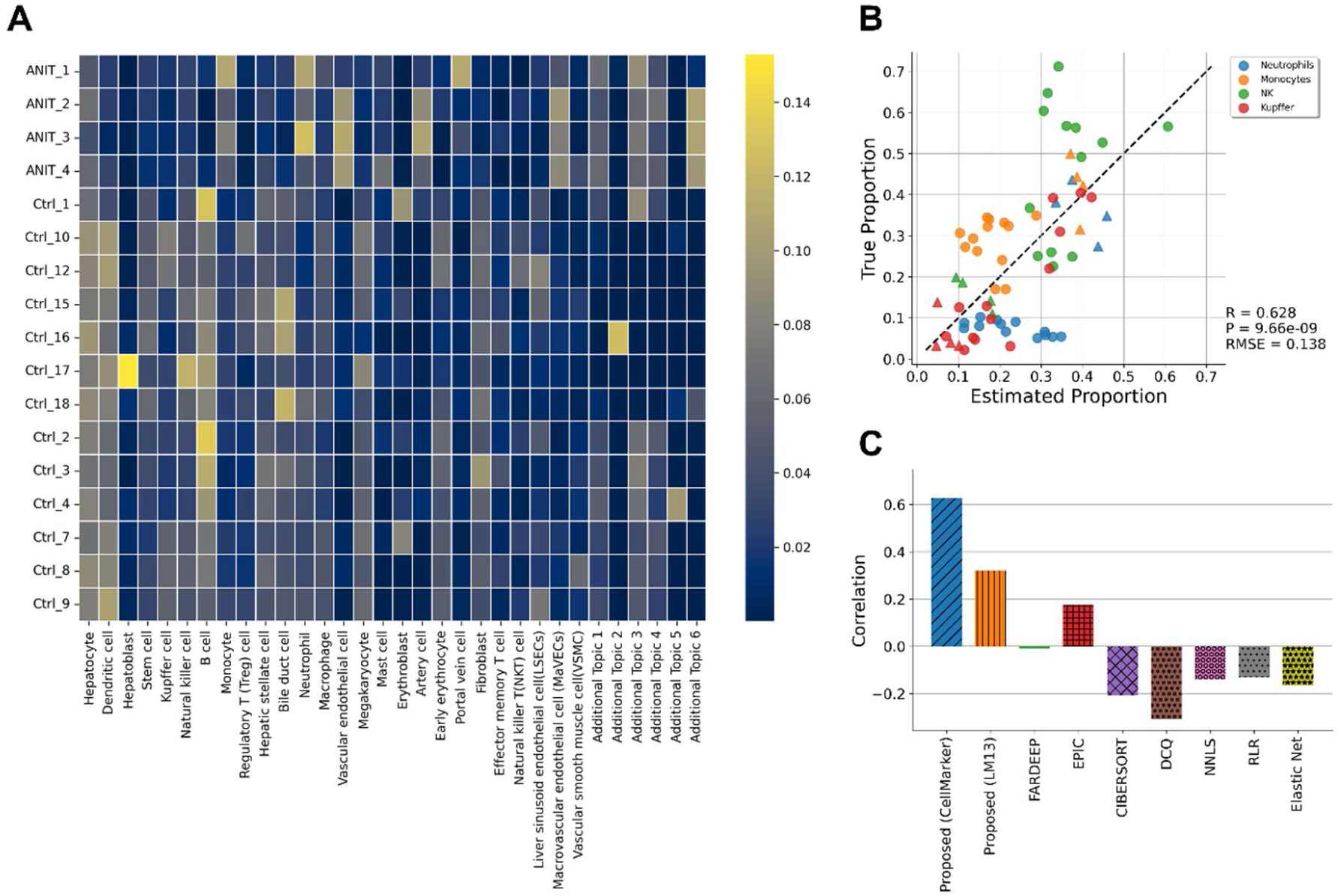
Prediction of immune cell trafficking in mouse liver tissue perturbed by alpha-naphthyl isothiocyanate (ANIT). **(A)** Comprehensive cell type proportions estimation using marker gene names collected in data-driven manner. **(B)** Scatterplot showing the estimated proportions of immune cells vs. measured values in the same dataset. Circles indicate control samples and triangles indicate samples treated with ANIT. **(C)** Performance comparison based on Pearson correlation with other notable deconvolution methods.

**Figure S12.**
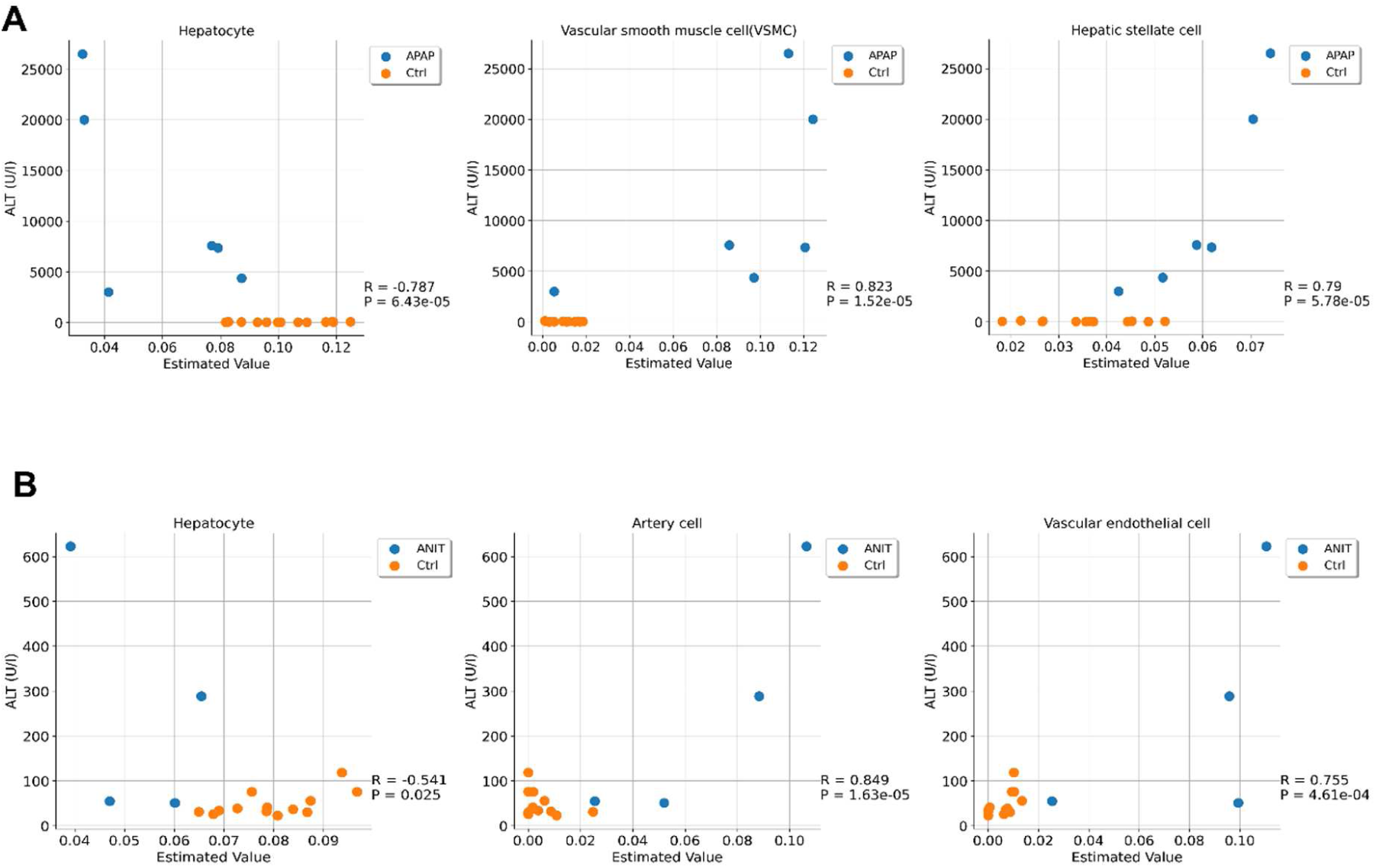
Relationship between the cell ratio estimated by GLDADec and the degree of injury. Scatterplots comparing estimated proportions and blood biochemistry value in **(A)** APAP and **(B)** ANIT treatment group, respectively. Alanine aminotransferase (ALT) is the most common marker of liver damage. High ALT values are a sign of severe liver damage.

**Figure S13.**
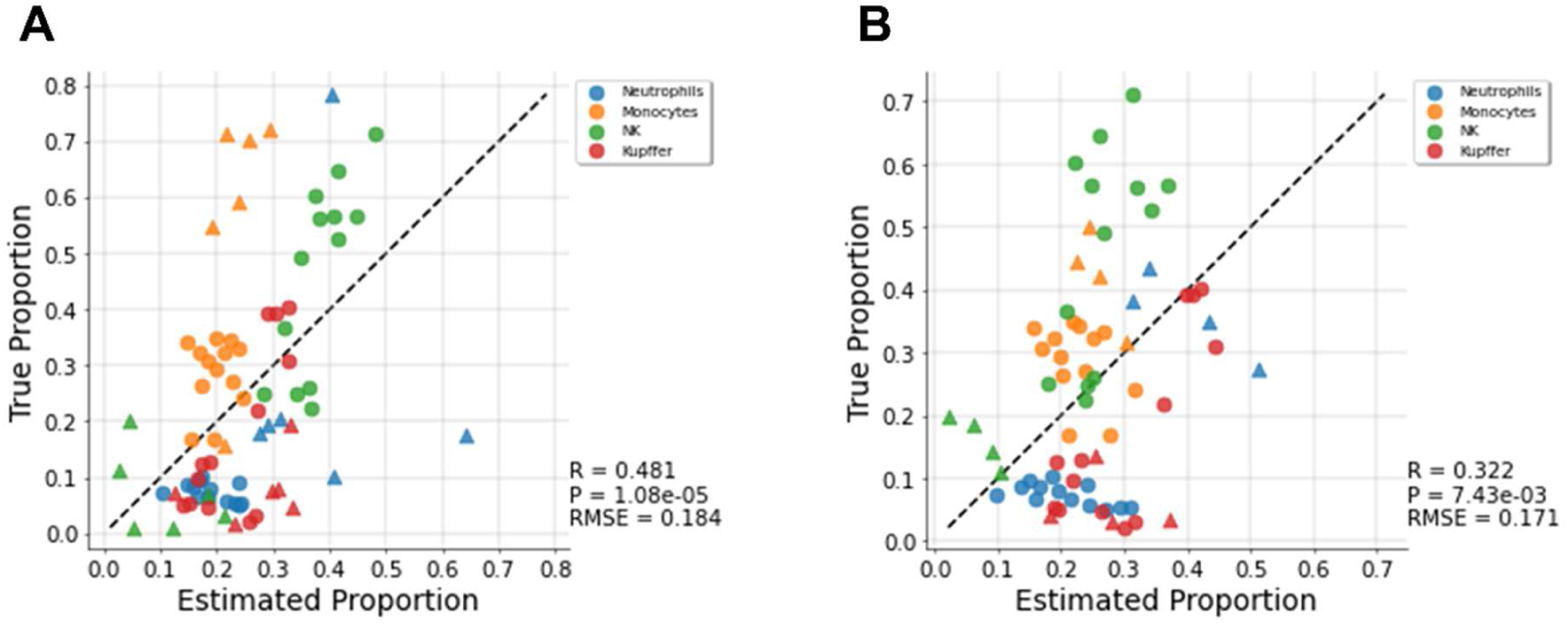
Estimation performance of GLDADec when using differentially expressed genes (DEGs) derived marker genes as prior information. Scatterplot showing the estimated proportions of immune cells vs. measured values for **(A)** acetaminophen and **(B)** alpha-naphthyl isothiocyanate administration group, respectively.

**Figure S14.**
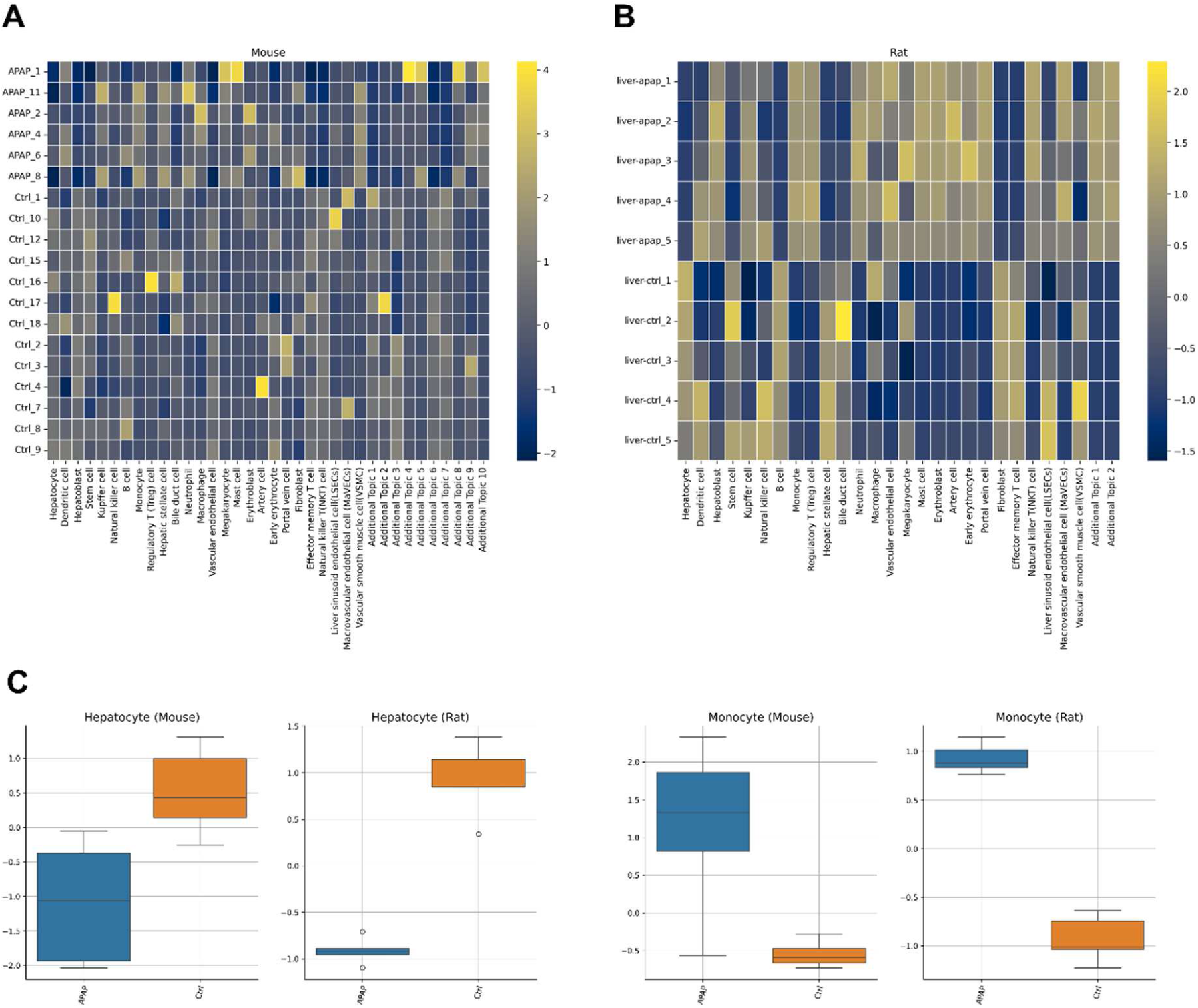
Comparison of immune cell trafficking patterns in mice and rats after acetaminophen (APAP) administration. Heatmaps showing the estimated comprehensive cell type proportions in **(A)** mice and **(B)** rats. **(C)** Common cell trafficking patterns between mice and rats.

**Figure S15.**
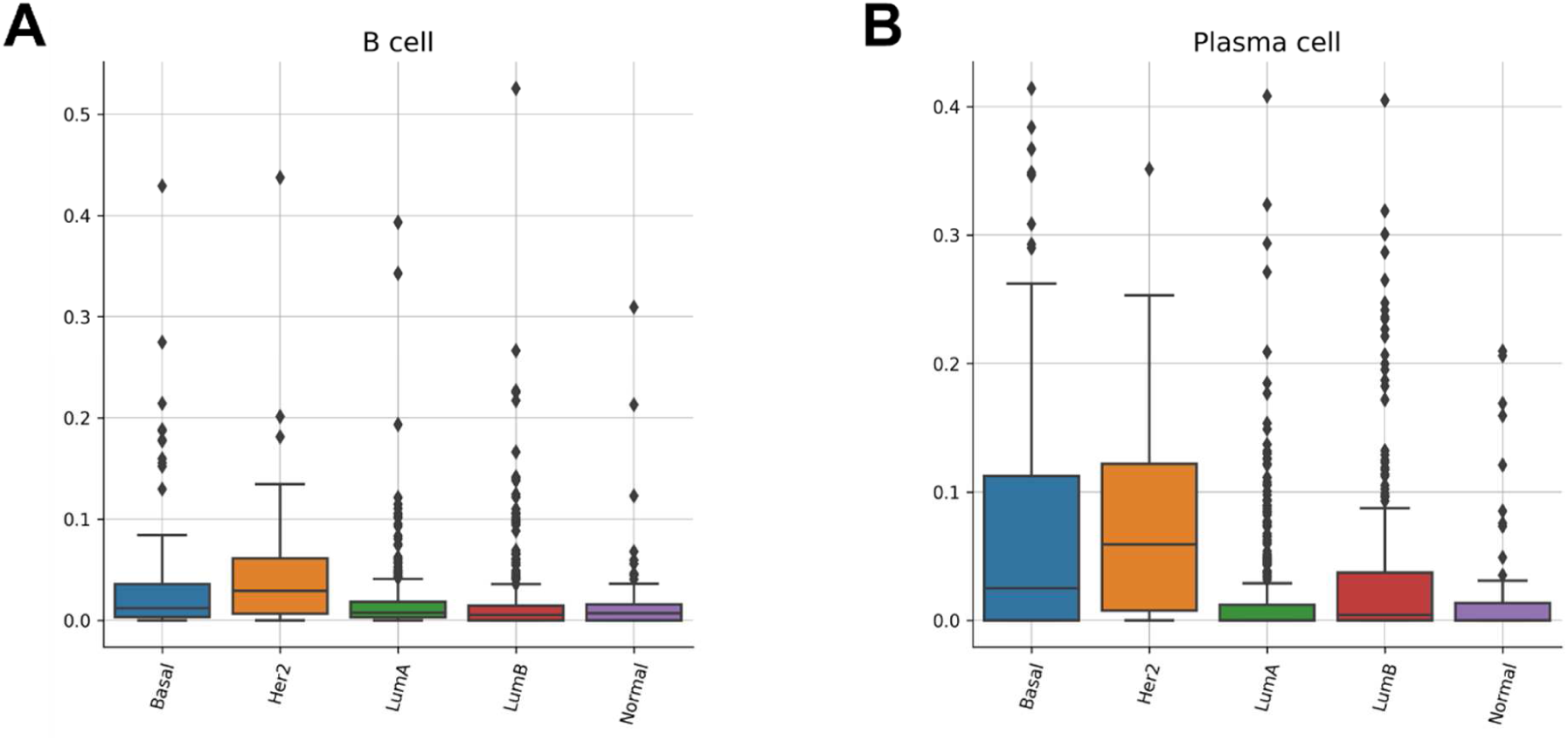
Boxplots showing specific accumulation and infiltration between each subtype of BRCA. Her2 subtype-specific accumulation was observed in **(A)** B cells and **(B)** plasma cells, respectively.

**Figure S16.**
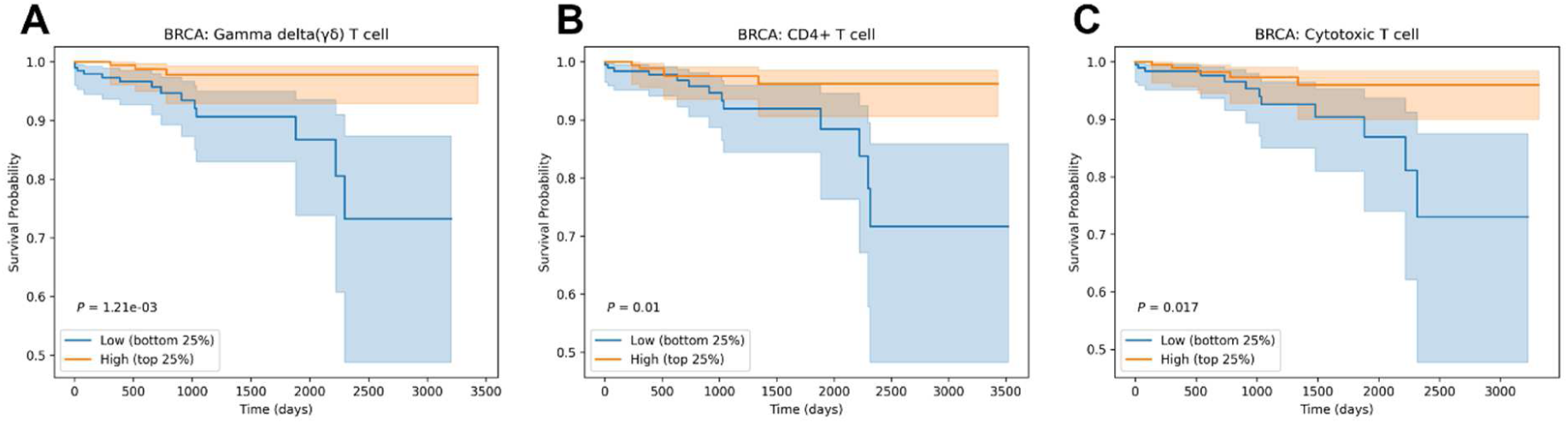
Survival analysis when samples are stratified by the accumulation of cells estimated by GLDADec. Kaplan-Meier plots showing survival associations with infiltration of **(A)** gamma delta (γδT) cells, **(B)** CD4+ T cells, and **(C)** Cytotoxic T cells in BRCA. Patients with the top 25^th^ percentile of the target cells were compared with those with the bottom 25^th^ percentile.

**Figure S17.**
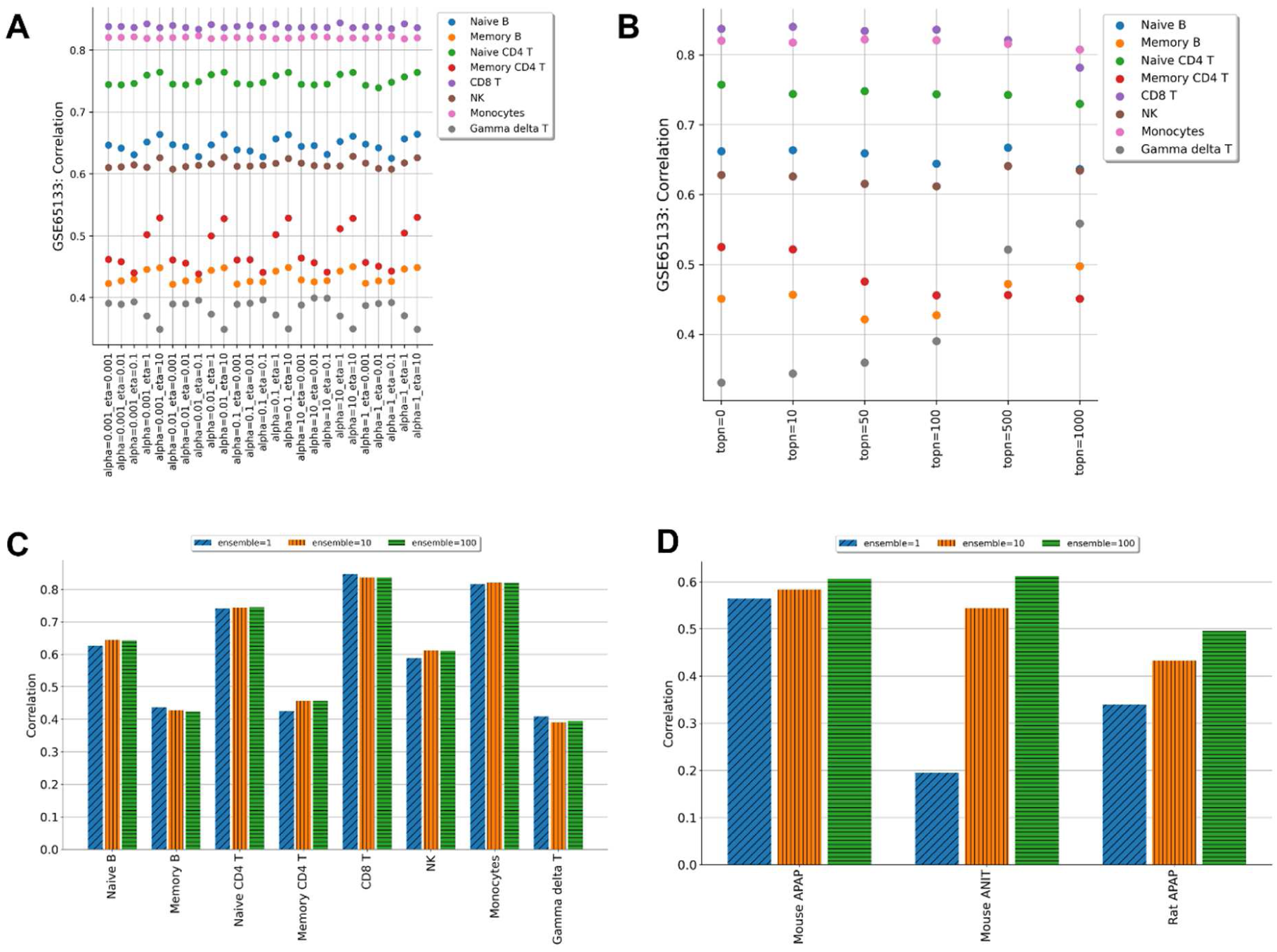
Hyperparameter sensitivity analysis. **(A)** Effect of the combination of hyperparameters α and η on the Dirichlet distributions. **(B)** Relationship between the number of additional genes with large coefficients of variation and estimation performance. Bar plots representing the contribution of the ensemble to improved estimation performance for **(C)** blood-derived samples and **(D)** tissue samples, respectively.

**Figure S18.**
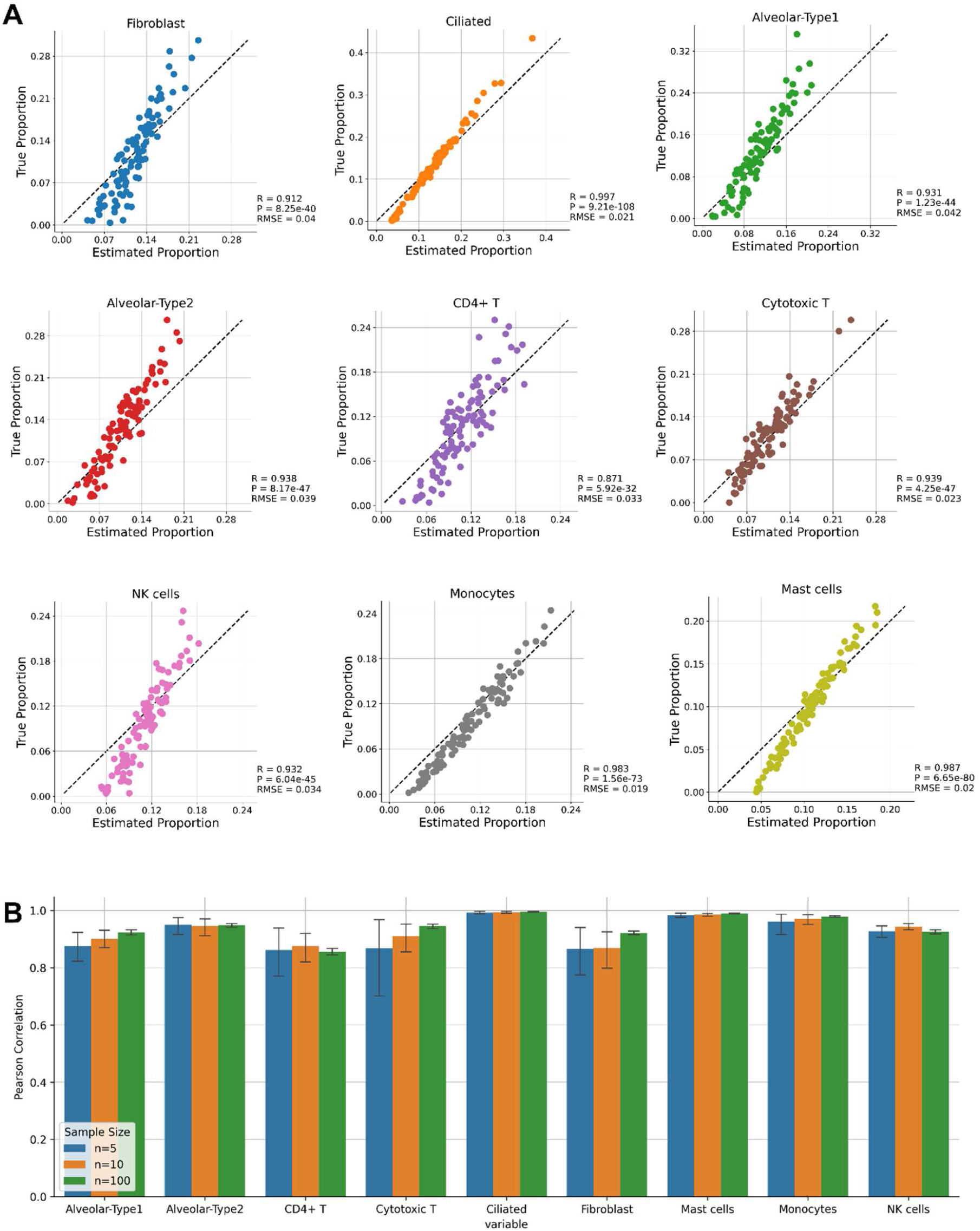
Assessing the robustness of the estimation performance of GLDADec. (A) Scatterplots showing estimated and ground-truth proportion for pseudo lung tissue data generated from single cell RNA-Seq. We generated 100 samples with randomly assigned proportions for the nine cell types. (B) Barplots showing the change in estimation performance with decreasing the number of samples.

**Figure S19.**
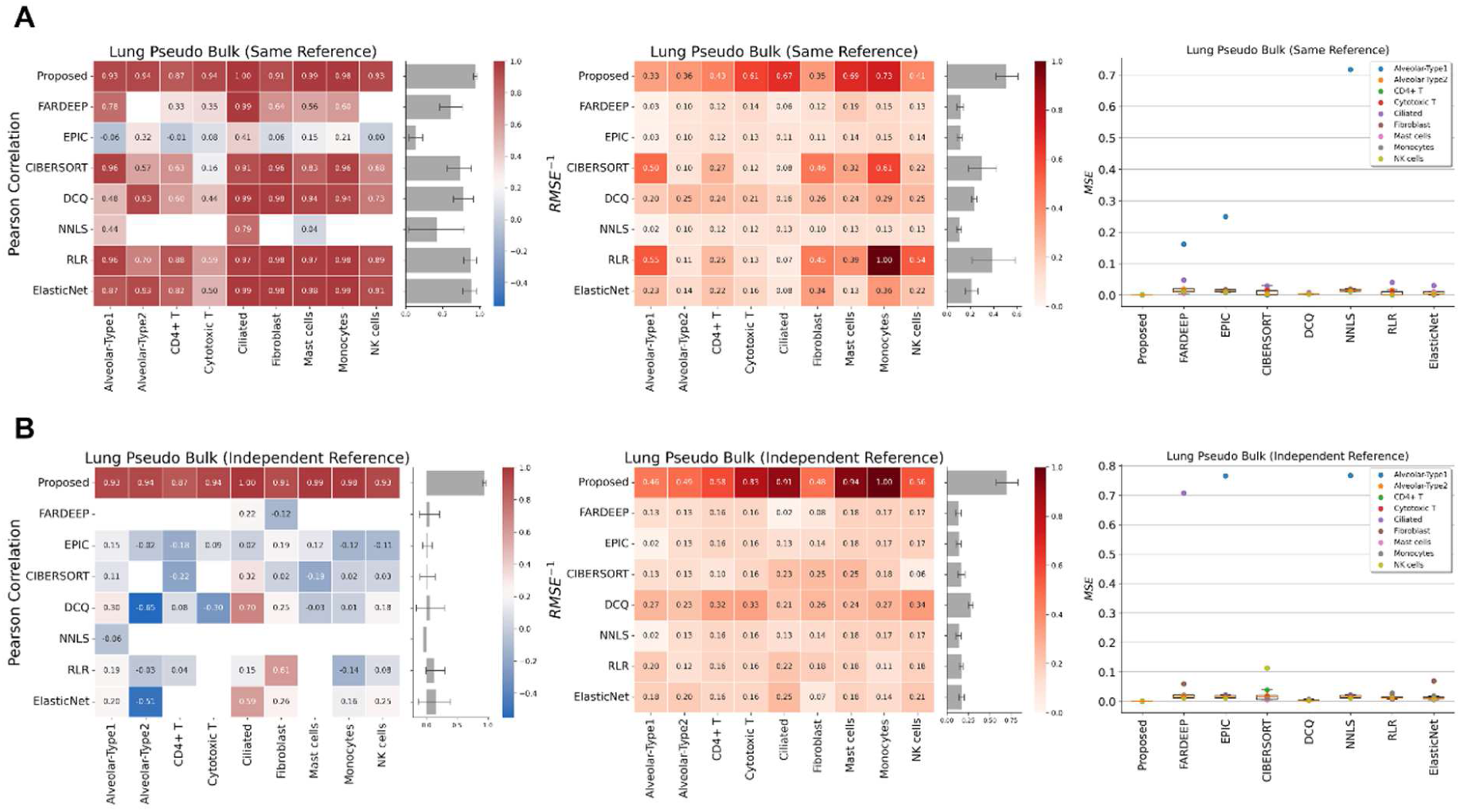
Assessing the robustness of GLDADec that is independent of the distribution of reference expression levels. Heatmap showing estimation performance when using a reference from (A) the same dataset and (B) an independent dataset. The left and center blocks show the Pearson correlation and the inversed root mean square error (RMSE) values (scaled 0 to 1), respectively. The barplots on the right shows the performance of each method across all cell types. The right block shows boxplots of mean square error (MSE). Each box extends from the 25^th^ percentile (bottom) to 75^th^ percentile (top), and the whisker indicates the farthest data point within 1.5-fold of inter-quartile range.

**Table S1.** Summary of datasets used in this study.

| Dataset | PMID | Size | Species | Source |
| --- | --- | --- | --- | --- |
| GSE65133 | 25822800 | 20 | Human | PBMC |
| GSE107572 | 31126321 | 9 | Human | PBMC/PMN |
| GSE60424 | 25314013 | 5 | Human | Whole blood |
| SDY67 | 27031986 | 267 | Human | PBMC |
| GSE107011 | 30726743 | 12 | Human | PBMC |
| ROSMAP | 32804935 | 41 | Human | Brain |
| GSE237801-APAP | 38187088 | 19 | Mouse | Liver |
| GSE237801-ANIT | 38187088 | 17 | Mouse | Liver |
| GSE239996-APAP | 37941435 | 10 | Rat | Liver |
| TCGA-BRCA | - | 1052 | Human | Breast |
| TCGA-LUAD | - | 577 | Human | Lung |
| TCGA-LIHC | - | 408 | Human | Liver |
| Tissue Stability Cell Atlas | 31892341 | - | Human | Lung (single cell) |
| GSE131907 | 32385277 | - | Human | Lung (single cell) |

**Table S2.** Gene Ontology (GO) enrichment analysis for the added topic that reconstructing missing neuron. The top 10 significantly enriched GO terms are shown with Benjamini–Hochberg adjusted p-values.

| Rank | GO Term | Adjusted P-value | Overlap |
| --- | --- | --- | --- |
| 1 | Chemical Synaptic Transmission (GO:0007268) | 0.001153 | {'GABRA1', 'SST', 'SNAP25', 'GAD2', 'GABRD', 'SLC1A1', 'SLC17A7', 'GAD1', 'SLC17A6'} |
| 2 | L-glutamate Import (GO:0051938) | 0.009144 | {'SLC17A6', 'SLC1A1', 'SLC17A7'} |
| 3 | Neurotransmitter Transport (GO:0006836) | 0.01298 | {'SNAP25', 'SLC17A6', 'SLC1A1', 'SLC17A7'} |
| 4 | Cellular Response To Nerve Growth Factor Stimulus (GO:1990090) | 0.01313 | {'MAPT', 'STMN2', 'SH3GL2'} |
| 5 | Positive Regulation Of Microtubule Polymerization Or Depolymerization (GO:0031112) | 0.01362 | {'MAPT', 'STMN2', 'PAK1'} |
| 6 | Glutamate Catabolic Process (GO:0006538) | 0.01362 | {'GAD2', 'GAD1'} |
| 7 | Gamma-Aminobutyric Acid Metabolic Process (GO:0009448) | 0.01362 | {'GAD2', 'GAD1'} |
| 8 | Anterograde Trans-Synaptic Signaling (GO:0098916) | 0.01362 | {'SNAP25', 'SST', 'GAD2', 'GABRD', 'SLC1A1', 'GAD1'} |
| 9 | Sodium-Dependent Phosphate Transport (GO:0044341) | 0.02114 | {'SLC17A6', 'SLC17A7'} |
| 10 | Regulation Of Mesenchymal Stem Cell Differentiation (GO:2000739) | 0.02114 | {'PDGFRA', 'SOX6'} |

**Table S3.**
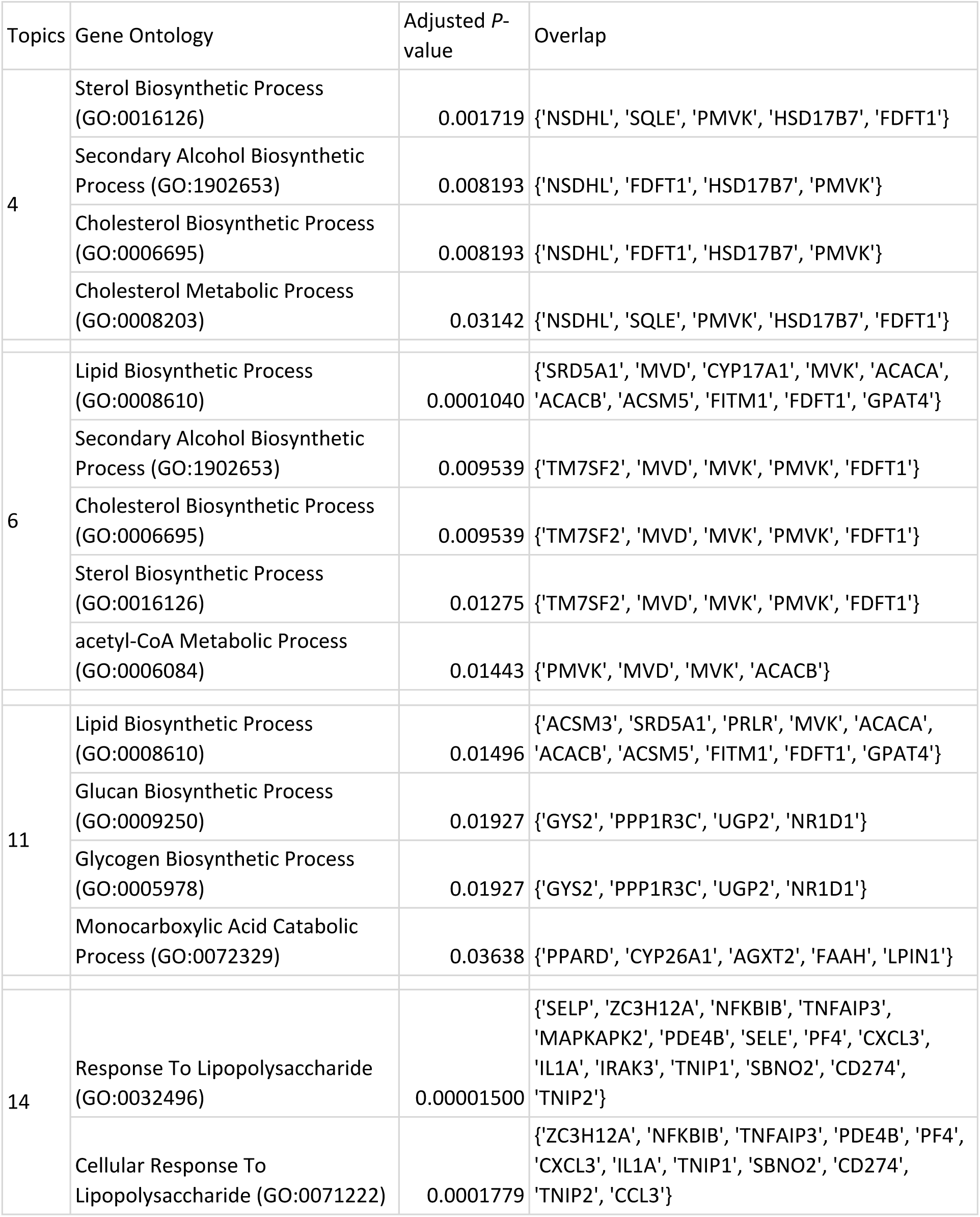
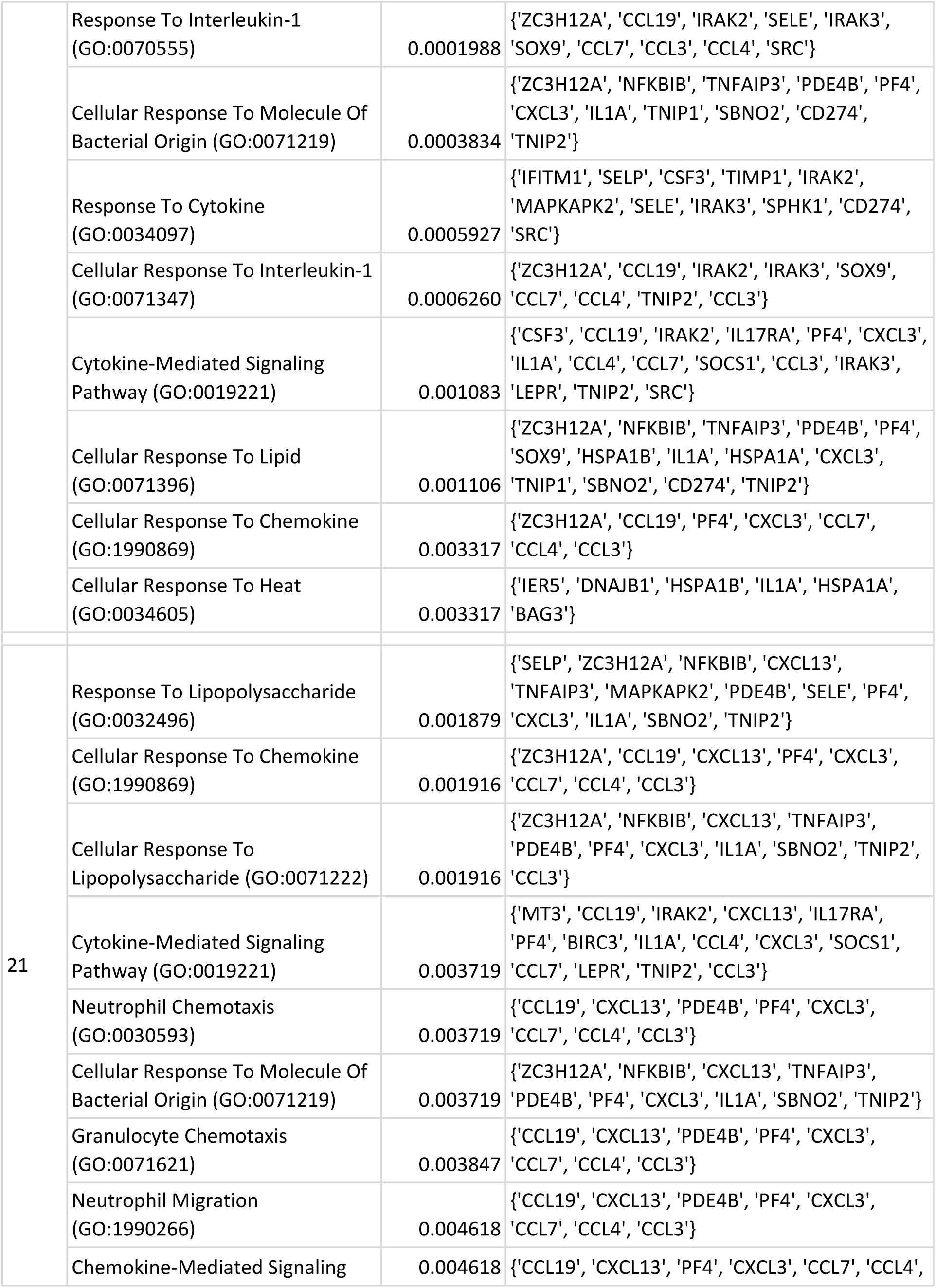
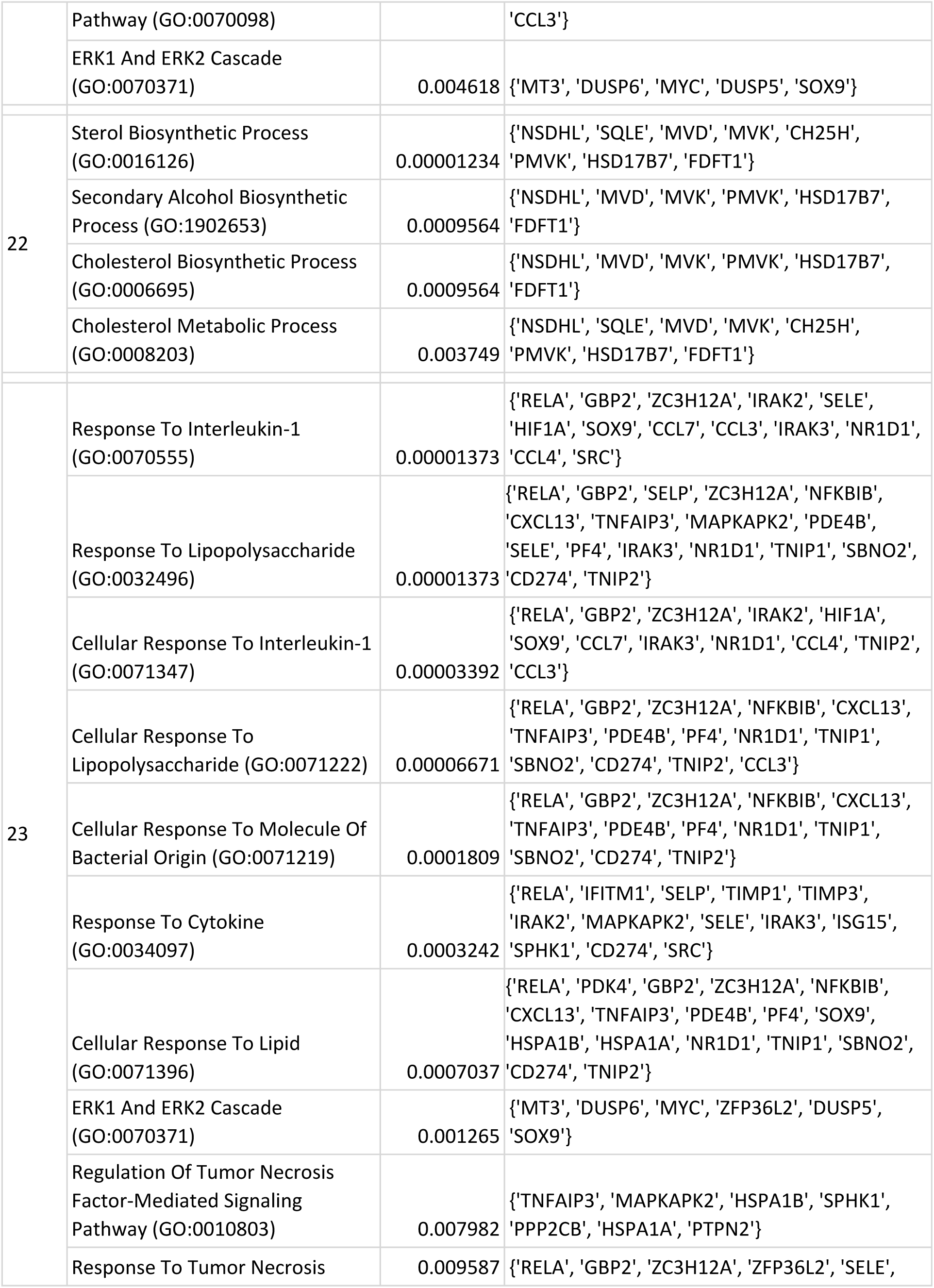
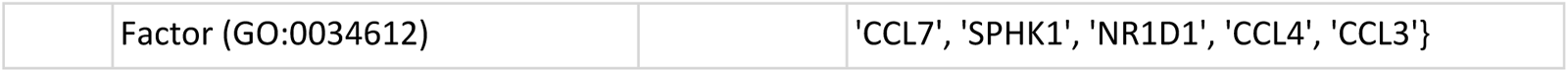
Gene ontology (GO) enrichment analysis. The top 10 significantly enriched GO terms are shown with Benjamini–Hochberg adjusted *P*-values.

**Table S4.**
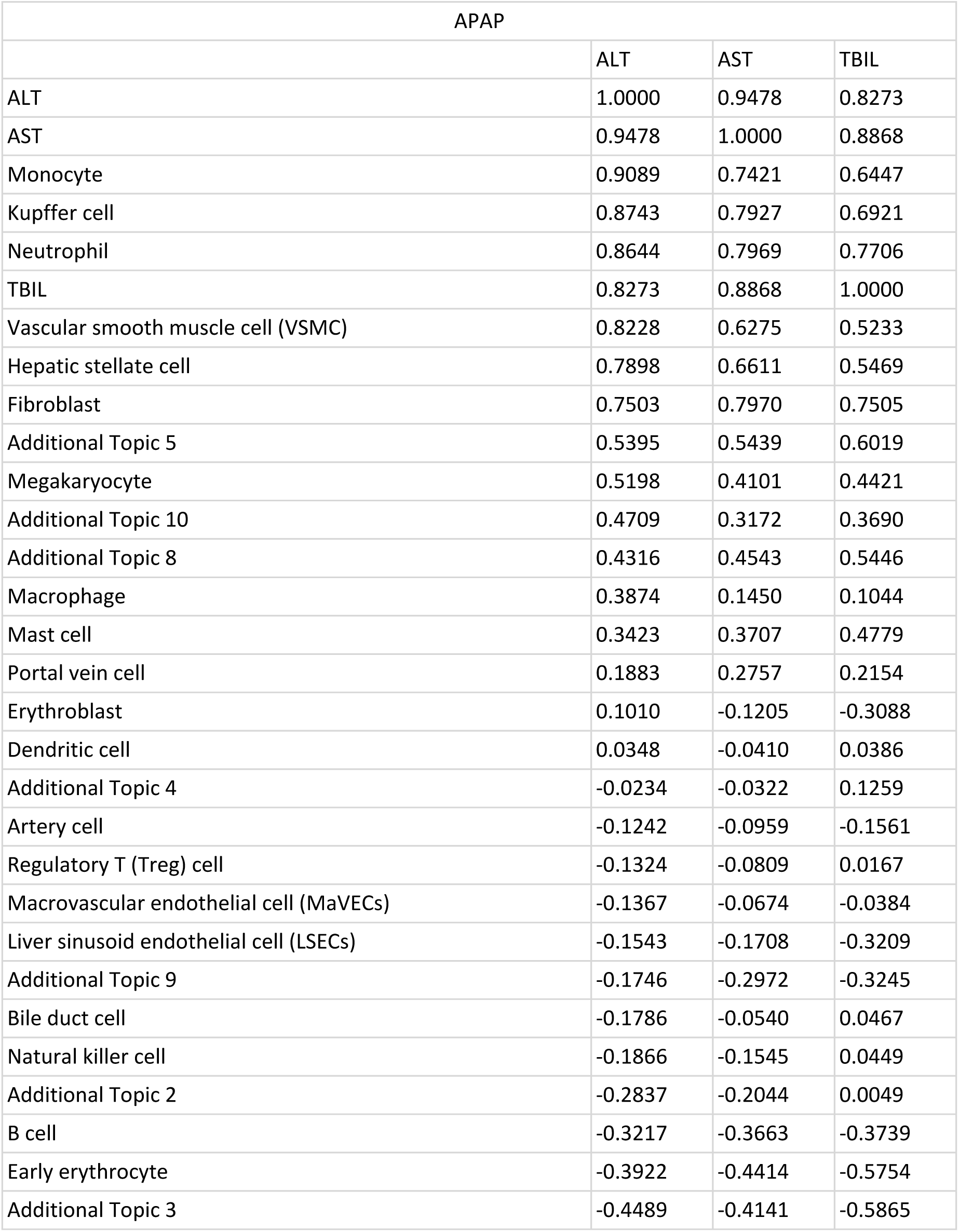
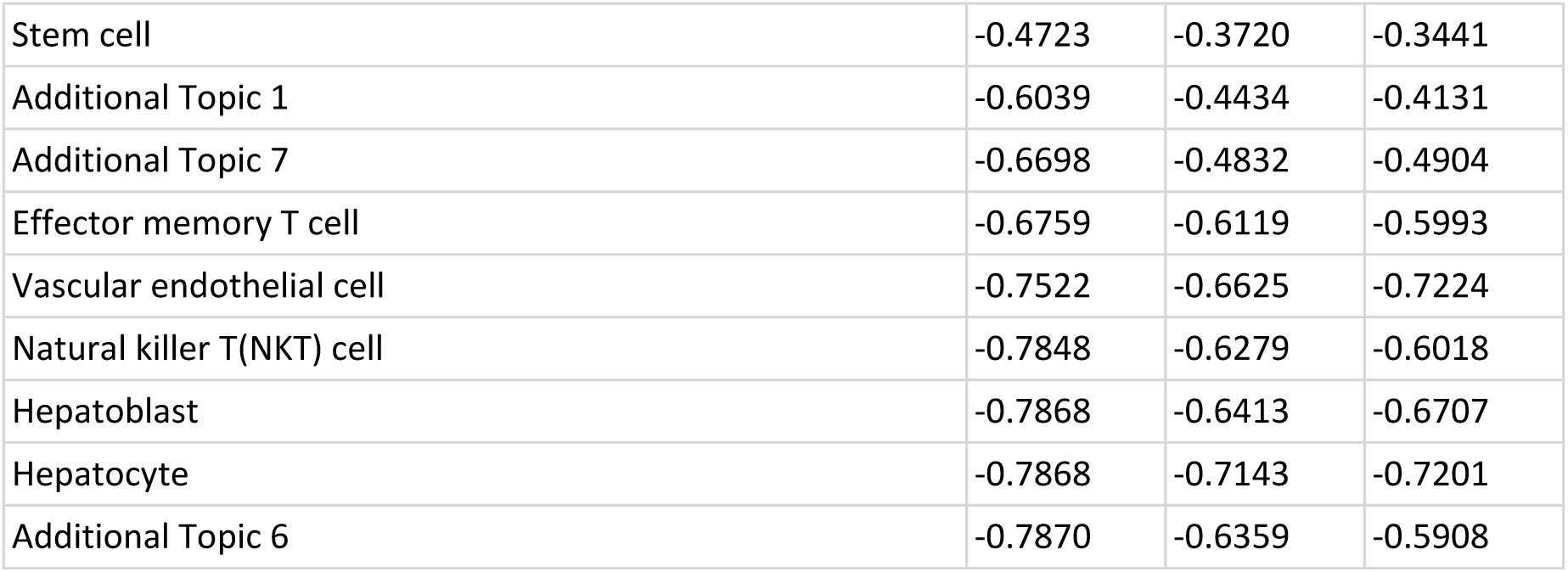
Pearson correlation between cell proportions estimated by GLDADec and various liver injury markers such as ALT, AST, and TBIL after acetaminophen (APAP) administration.

**Table S5.** Pearson correlation between cell proportions estimated by GLDADec and various liver injury markers such as ALT, AST, and TBIL after alpha-naphthyl isothiocyanate (ANIT) administration.

| ANIT |  |  |  |
| --- | --- | --- | --- |
|  | ALT | AST | TBIL |
| ALT | 1.0000 | 0.9837 | 0.8807 |
| AST | 0.9837 | 1.0000 | 0.9075 |
| TBIL | 0.8807 | 0.9075 | 1.0000 |
| Artery cell | 0.8490 | 0.8253 | 0.8658 |
| Vascular endothelial cell | 0.7549 | 0.7216 | 0.8763 |
| Additional Topic 6 | 0.7216 | 0.6840 | 0.8239 |
| Neutrophil | 0.6691 | 0.7145 | 0.7333 |
| Macrovascular endothelial cell (MaVECs) | 0.6394 | 0.5930 | 0.7701 |
| Monocyte | 0.4994 | 0.5086 | 0.6127 |
| Additional Topic 4 | 0.2371 | 0.2374 | 0.3291 |
| Additional Topic 1 | 0.2226 | 0.2758 | 0.4081 |
| Mast cell | 0.2084 | 0.1795 | 0.4790 |
| Portal vein cell | 0.2069 | 0.2387 | 0.3103 |
| Additional Topic 3 | 0.1772 | 0.2330 | 0.2685 |
| Effector memory T cell | -0.0085 | 0.0074 | -0.0951 |
| Additional Topic 2 | -0.0376 | -0.0676 | -0.0619 |
| Hepatoblast | -0.0565 | -0.0824 | -0.0485 |
| Natural killer T(NKT) cell | -0.0570 | -0.1021 | -0.2580 |
| Additional Topic 5 | -0.1332 | -0.1093 | -0.1711 |
| Erythroblast | -0.2500 | -0.1994 | -0.2383 |
| Natural killer cell | -0.2920 | -0.3137 | -0.3087 |
| Regulatory T (Treg) cell | -0.3018 | -0.3478 | -0.4882 |
| Kupffer cell | -0.3059 | -0.3175 | -0.4427 |
| Vascular smooth muscle cell (VSMC) | -0.3061 | -0.2617 | -0.1206 |
| Stem cell | -0.3683 | -0.4029 | -0.5304 |
| Liver sinusoid endothelial cell (LSECs) | -0.3801 | -0.3444 | -0.4674 |
| Bile duct cell | -0.4157 | -0.4109 | -0.4434 |
| Macrophage | -0.4244 | -0.4285 | -0.4181 |
| Early erythrocyte | -0.4520 | -0.4542 | -0.6601 |
| Megakaryocyte | -0.4991 | -0.4974 | -0.5964 |
| Fibroblast | -0.5285 | -0.5247 | -0.6216 |
| Hepatocyte | -0.5410 | -0.5995 | -0.7191 |
| Dendritic cell | -0.5531 | -0.5706 | -0.6903 |
| B cell | -0.5755 | -0.5371 | -0.6592 |
| Hepatic stellate cell | -0.5780 | -0.5506 | -0.6094 |

## Supplementary notes

### Note S1. Data preparation and processing

In this section, we will outline the detailed data preparation and processing methods. The processed data can be accessed on our Github repository (https://github.com/mizuno-group/GLDADec).

#### Blood-derived benchmarking data

The PBMC microarray, PBMC/PMN RNA-Seq, and whole blood RNA-Seq datasets were obtained from the NCBI Gene Expression Omnibus (GEO), with accessions GSE65133, GSE107572, and GSE60424 respectively as processed series matrix files [1–4]. These blood-derived samples are commonly utilized to evaluate deconvolution methods [1–3,5–7]. In addition, we obtained PBMC RNA-Seq data with known more detailed cell type classification, SDY67 and GSE107011 [4,8]. For the SDY67 dataset, we obtained it from Immport database [8]. These datasets were available in the study by Swapna et al [9]. All these datasets were derived from human samples, and the transcript IDs were converted to HGNC symbols using files available from Biomart [10]. Note that only immune cell types measured by flow cytometry and included in the benchmark data were selected for evaluation.

A list of typical marker genes for the cells measured in the benchmark datasets was established using domain knowledge and is provided in Supplementary File S1.

#### Brain dataset for benchmarking

We acquired bulk data for the prefrontal cortex from the Religious Orders Study / Memory and Aging Project (ROSMAP) for brain tissue analysis [11]. This dataset offers cell type proportions determined through immunohistochemistry, serving as the ground truth. The processed gene expression matrix were obtained from GitHub repository (https://github.com/ellispatrick/CortexCellDeconv) reported in the original study, and the ground-truth proportions are reported by Swapna et al [9]. Samples that retain ground-truth information ware selected to evaluate estimation performance.

Marker genes for each cell types were defined based on domain knowledge. Supplementary File S2 contains a list of these marker genes.

#### Tissue data with perturbation

To assess the estimation of immune cell trafficking in tissues, we retrieved bulk RNA-Seq data from mice and rats with accessions GSE237801 and GSE239996. These data encompass the tissue transcriptome from drug-induced liver injury models and the principal immune cell proportions in the tissues.

Transcripts per kilobase million (TPM) values were obtained for the data, and the transcript IDs were converted to gene symbols. The median values were selected for genes with duplicated names. The control samples and the samples of interest, such as the acetaminophen-treated group, were merged and used as targets to estimate the cellular changes associated with the perturbation. To highlight the differences in the trafficking of immune cells due to perturbation, a feature-wise minmax correction was performed as a preprocessing step to emphasize differences in expression levels between samples.

Cell types verified by flow cytometry were evaluated for deconvolution, including neutrophils, monocytes, NK cells, and Kupffer cells in mice, and B cells, CD4+ T cells, CD8 + T cells, neutrophils, monocytes, and NK cells in rats.

Marker genes for cell types linked to mouse liver were obtained from CellMarker [12]. Cleansing was performed by integrating homologous cell names and eliminating abstract cell names, and finally marker genes specific to the 34 cell types were defined (Supplementary File S3). As there is no established database for marker gene names in rats, we obtained classical cell-type markers that are highly expressed in rats, mice, and humans, as reported in the study by Natasha et al [13].

#### Clinical data

To assess the utility of GLDADec in the context of human clinical data, we obtained three distinct types of tumor datasets, comprising 1052 breast invasive carcinoma (BRCA) samples, 577 lung adenocarcinoma (LUAD) samples, and 408 liver hepatocellular carcinoma (LIHC) samples, from the GDC data portal on November 16, 2022 [14]. Marker genes for the cell types associated with each human tissue were obtained from the CellMarker database [12]. The expression were normalized using TPM, and quantile normalization was performed to mitigate the impact of the distribution of expression levels in each sample on the estimated values. Due to the large number of samples and the high computational demands of these datasets, the linear scale expression was transformed into a sparser matrix by dividing it by a constant value of 1000.

#### Pseudo lung bulk dataset

The single-cell transcriptome data for cells isolated from human lung tissue was sourced from the Tissue Stability Cell Atlas (TSCA) [15]. A pseudo-bulk dataset was generated comprising five immune cell types (CD4+ T cells, cytotoxic T cells, mast cells, monocytes, and NK cells) and four lung background cell types (alveolar type I, alveolar type II, ciliated cells, and fibroblasts). The generation process involved the following steps: (1) Determining 500 cells each for immune and background types, totaling 1000 cells. (2) Randomly assigning proportions to the constituent cell types under a sum-to-one constraint. (3) Partitioning the data into main and test sets, extracting the corresponding cell counts from the main data, and summing the count data. (4) Repeating this process to create a pseudo-bulk dataset with a chosen sample size. We computed the mean expression level of cells within the test group and established the reference using the same batch as the pseudo-bulk dataset. Additionally, we established a reference derived from a different batch, using accession GSE131907, an independent lung single-cell RNA-Seq dataset published by Kim et al [16].

For TSCA dataset, the raw data was downloaded in h5ad format and processed using the python scanpy library. Ensemble transcript IDs were converted to HGNC symbols using files available from Biomart [10]. For the other dataset, GSE131907, we downloaded raw matrix with Unique molecular identifier and annotation file, respectively. Since this is a txt formant with a huge number of lines, we read each line (barcode) and extracted the cell types of interest for downstream task. This analysis flow is also available on our GitHub repository (https://github.com/mizuno-group/GLDADec).

### Note S2. Condition of experiments

In this section, we provide a comprehensive description of the execution environment of the proposed and existing methods in various scenarios, which can be replicated in our GitHub repository (https://github.com/mizuno-group/GLDADec).

#### Implementation and data collection of other deconvolution methods

This subsection describes the common setting to perform all the benchmarking, including implementation and hyperparameters.

FARDEEP, EPIC, CIBERSORT, DCQ, NNLS, RLR, and Elastic Net are bulk reference deconvolution methods [1,7,17,18]. For FARDEEP, we used the corresponding R package (https://cran.r-project.org/web/packages/FARDEEP/index.html) with the default parameter settings to obtain the estimated values of absolute abundance of cells. For EPIC, we employed the online portal (https://epic.gfellerlab.org/) to get cell fractions following the default settings. The CIBERSORT website has now been incorporated into the CIBERSORTx website, and we utilized its online portal (https://cibersortx.stanford.edu/). All parameters, including the disabling of quantile normalization, were kept at default values. For DCQ, we used the R package (https://cran.r-project.org/web/packages/ADAPTS/index.html) implemented by Samuel et al. with the default parameter settings [19]. For the remaining NNLS, RLR, and Elastic Net, we employed the code from the GitHub repository (https://github.com/mizuno-group/LiverDeconv) implemented in our previous study with default parameter settings.

For more detailed cell type estimation benchmarking tasks, five state-of-the-art methods leveraging single-cell RNA-Seq or marker gene information were used, including GTM-decon, BayesPrisim, CIBERSORTx, MuSiC, and BSEQ-sc [9,20–23]. GTM-decon offeres three different gene selection methods. We obtained results with the *hvg* method, which selects highly variable genes, a scenario like our method. Additionally, it is known that GTM-decon achieves better performance using multiple topics per cell type, and the reported results were obtained under conditions with five topics per cell type. For BaysPrism, the metadata column for tumor status set to 0 for all cell types, and the online web portal (https://www.bayesprism.org/) was utilized. For CIBERSORTx, the settings were identical to those used for CIBERSORT as described above. For the remaining methods, MuSiC and BSEQ-sc, the reported results were obtained using recommended or default settings. Note that the baseline scores for these five state-of-the-art methods were reported in a recent study by Swapna et al. (https://github.com/li-lab-mcgill/gtm-decon), and some parameter settings were confirmed by contacting the authors.

#### Evaluation metrics for deconvolution

We evaluated the performance using several metrics, including Pearson correlation, rooted mean square error (RMSE), and mean square error (MSE). As a preprocessing step, we adjusted the sum of the proportions of cell types to 1 for both the deconvolution output and the ground-truth. Note that the cell types subject to adjustment are those common between the output and the ground-truth data. In other words, the cell type range constrained by sum to 1 was identical, ensuring comparable evaluations.

#### Benchmarking with human blood samples

We employed a linear scaled blood-derived benchmark dataset, utilizing marker gene names defined by domain knowledge (see Supplementary File S1) as partial prior information. Genes exhibiting outliers greater than 2σ from the log-normal distribution of gene expression levels were eliminated for each sample, and 100 genes displaying substantial sample-wide coefficients of variation (CV) were incorporated into the analysis. This number of genes to be added serves as a hyperparameter and is further discussed in the main text. It is important to note that the benchmark data originate from blood samples, and no additional topics were integrated into the model since the constituent cell types are clearly delineated. To correct for the expression level on a linear scale, the values were divided by constants specific to each dataset, i.e., 10, 200, 100, 1000, and 10 for GSE65133, GSE107572, GSE60424, SDY67, and GSE107011, respectively. This operation was implemented to save computational resources and does not alter the relative gene expression levels of each sample.

We conducted a comparison of the performance of GLDADec with seven alternative deconvolution methods, including FARDEEP, EPIC, CIBERSORT, DCQ, NNLS, RLR, and Elastic Net [1,7,17,18]. These methods are all reference-based and require prior knowledge of a gene signature matrix as the reference. The LM22 definition provided by Newman et al. is commonly used as a reference for human blood-derived samples and was also obtained from their work and utilized in our analysis [1]. Notably, LM22 was defined in the same study that generated GSE65133, one of the benchmark datasets used in our analysis, making it a compatible reference. The competing methods have diverse development backgrounds and accept different formats of expression data as input. We adhered to their default settings whenever possible. Specifically, we employed log-normalized gene expression matrices for DCQ and FARDEEP, and a non-log linear scaled matrix for EPIC and CIBERSORT. The remaining NNLS, RLR, and Elastic Net did not specify the input format, so we adapted them to match the scale of the reference data. To eliminate the influence of differences in scale between the analysis object and the reference on the results, we converted LM22 to a log-scale in the DCQ and FARDEEP analysis.

We next compared GLDADec with five state-of-the-art methods, including GTM-decon, BayesPrisim, CIBERSORTx, MuSiC, and BSEQ-sc [9,20–23]. Baseline scores for these methods are reported in recent study by Swapna et al [9]. We downloaded the estimated values for each method and the ground-truth information of cell type proportion from “fig2_data.tar.gz” file stored in GTM-decon’s GitHub repository (https://github.com/li-lab-mcgill/gtm-decon). In the original benchmark, dendritic cells (DC) and plasmacytoid dendritic cells (pDC) were distinguished, but in this study, pDCs were assumed to be included in DC, and these two cell estimates were combined.

#### Comprehensive cell type analysis for mouse perturbed tissue data

The dataset GSE237801 comprises bulk RNA-Seq data from mouse liver tissue under various compound administration and immune cell proportions measured by flow cytometry, which is a suitable benchmark dataset for deconvolution methods for tissues [24]. This study primarily focused on the acetaminophen (APAP) and alpha-naphthyl isothiocyanate (ANIT) treatment groups, examining the performance of deconvolution methods in detecting perturbation-induced changes in immune cell proportions relative to control samples. It is important to note that the tissue sample is a heterogeneous cell population, and subsequent analyses incorporate additional modeling considerations.

In our analysis of benchmarking using human blood datasets, GLDADec utilized linear scale expression levels as input and marker genes as prior information. In the examination of tissues, it is anticipated that transcriptome variation will arise due to the presence of unknown cells and confounding factors. To address this, we gathered liver-related cell types and their associated marker genes in a data-driven manner from the CellMarker database [12]. Moreover, to account for the diverse factors affecting the tissue transcriptome, we included 1000 genes with substantial coefficients of variation, which adequately describe the state of the tissue.

We contrasted the performance of GLDADec with seven existing reference-based methods, which are identical to those analyzed using the blood-derived benchmark dataset. Since the LM22 expression levels are derived from human samples, we employed the mouse-derived LM13, as previously defined in our analysis, as a reference to account for species differences in the mouse data. Additionally, we performed an analysis in which differentially expressed gene (DEGs) names derived from the same LM13 were considered as markers and subjected to GLDADec.

#### Application to rat data

GSE239996 contains data on rat liver tissue bulk RNA-Seq under various compound administration and immune cell proportions measured by flow cytometry [25]. In this study, we concentrated on the acetaminophen (APAP) treatment groups and assessed the efficacy of the deconvolution method in detecting immune cell proportion alterations induced by the compound compared to control samples. Similar to the analysis of the mouse data, 1000 genes with significant expression variation were included.

In this section, we defined three types of marker gene names for GLDADec analysis. The first set of classical cell type markers were collected from Figure 4A of Natasha’s work [13]. The second marker is the same as defined in the comprehensive analysis of mice from CellMarker. For the last set, DEGs were calculated for the six cell types (LM6) out of LM13, and the corresponding gene names were designated as markers. Utilizing LM6 as a reference, we compared its performance with seven competing methods.

#### Application to tumor samples

The TCGA RNA-seq data were obtained from numerous experiments and consist of a large number of samples, but also various confounding factors. To address this, we performed quantile normalization and aligned the distribution of each sample. However, the policy of eliminating genes with values that are more than two standard deviations away from the mean did not ensure enough genes, so we instead included genes with large expression variation in this analysis.

We gathered marker gene names from the background tissues of each cancer subtype using CellMarker and utilized this information as prior knowledge. Specifically, we obtained markers for 47, 73, and 80 cell types in a data-driven manner for breast, lung, and liver cancers, respectively. It is important to note that these markers were derived from human clinical data. Marker genes in human tissues have been well studied and are informative and comprehensive. Therefore, we performed modeling considering only one additional topic as “others”.

#### Application to pseudo lung bulk dataset

We included 100 genes with large coefficients of variation in the analysis, without considering additional topics. The setting is the same as for the analysis of blood samples, and this is since the cell types that comprise the sample are relatively explicit. In addition, the count data was corrected by dividing the count data by a constant of 100 to reduce the computational cost when GLDADec was performed.

### Note S3. Proof of 1.5

A Dirichlet distribution is defined with parameter α = (α_1_, α_2_, … , α_K_) as follows:

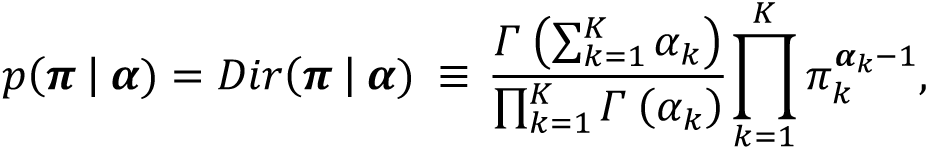

where, considering the graphical model of Latent Dirichlet Allocation (LDA), we obtain the following equation:

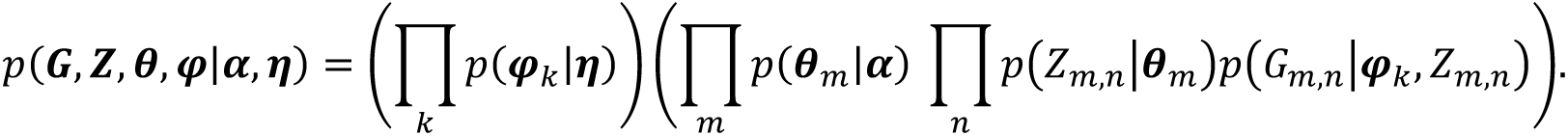

Now, to consider the joint distribution of p(G, Z | α, 1), both θ and φ were integrated out. Let *n_k,v_* = 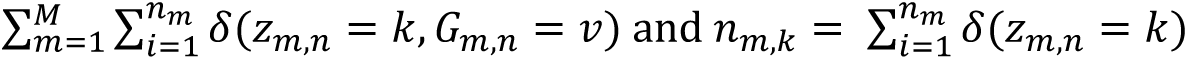, and the equation is written as follows:

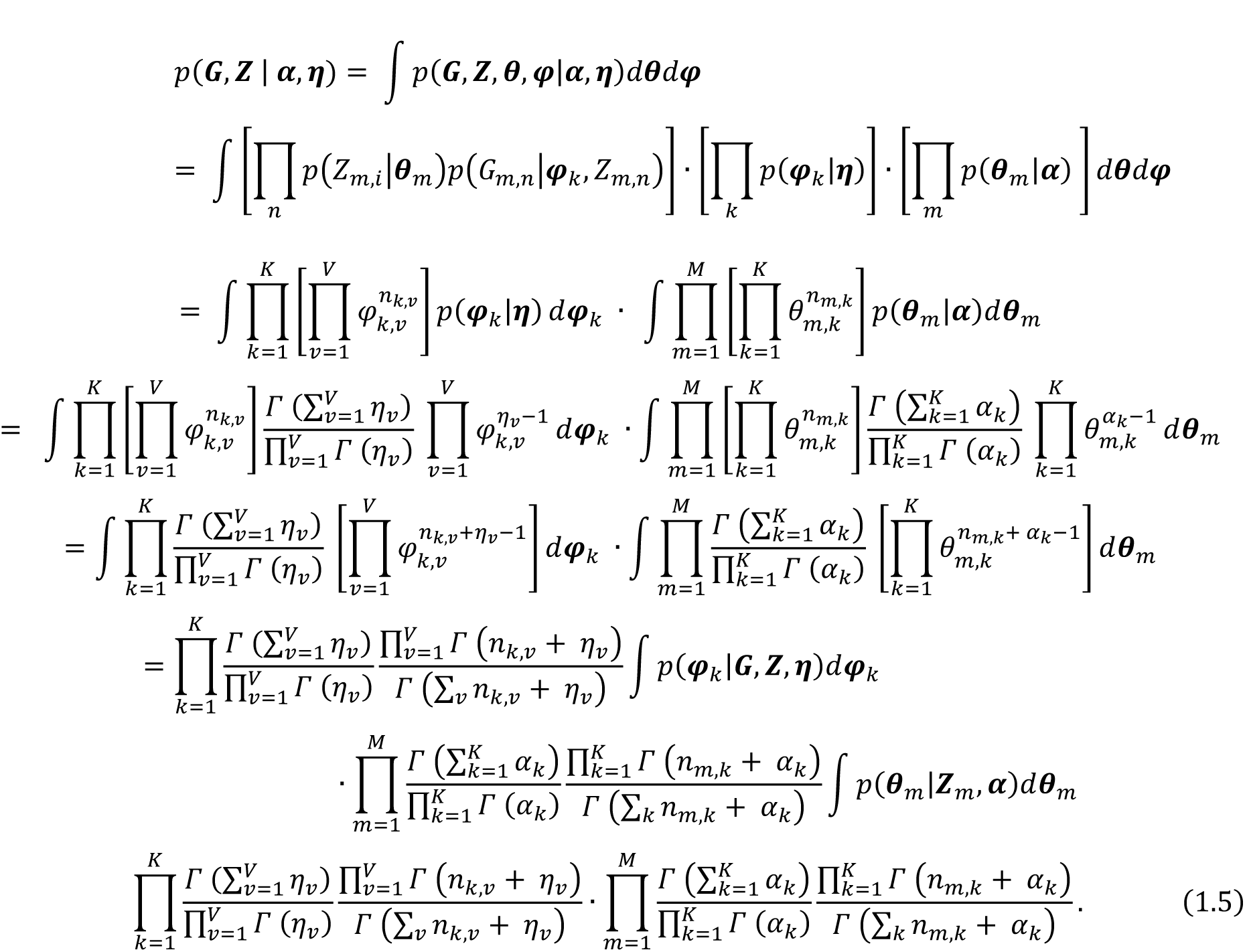

